# Increasing growth temperature alters the within-host competition of viral strains and influences virus genetic variation

**DOI:** 10.1101/2020.07.06.190173

**Authors:** Cristina Alcaide, Josep Sardanyés, Santiago F. Elena, Pedro Gómez

## Abstract

The emergence of viral diseases in plant crops hamper the sustainability of food production, and this may be boosted by global warming. Concurrently, mixed viral infections are becoming common in plants, of which epidemiology are unpredictable due to within-host virus-virus interactions. However, the extent in which the combined effect of variations in the abiotic components of the plant ecological niche (*e.g.*, temperature) and the prevalence of mixed infections (*i.e.*, within-host interactions among viruses) affect the evolutionary dynamics of viral populations is not well understood. Here, we explore the interplay between ecological and evolutionary factors during viral infections, and show that two individual strains of pepino mosaic virus (PepMV) coexisted in a temperature-dependent continuum between neutral and antagonistic interactions in tomato plants. After a long-term infection, the mutational analysis of the evolved viral genomes revealed strain-specific single-nucleotide polymorphisms that were modulated by the interaction between the type of infection and temperature. Mathematical modeling allowed us to asses a thermal reaction norm for both strains, which indicated that viral replication rates were increased along with increasing temperature in mixed infections, with a remarkable strain-dependent effect. These results suggest that the growth temperature is an ecological driver of virus-virus interactions, with an effect on the genetic diversity of individual viruses co-infecting a host. This research provides insights into the effect that climate change will have on the evolutionary dynamics of viral populations.

## INTRODUCTION

Emerging plant viral diseases are causing widespread epidemics in crops and represent a major burden to plant health with severe ecological, socio-economic, and political consequences^1,2^. Among many others, notable examples of viral diseases are tomato yellow leaf curl virus, which is devastating tomato production worldwide^3^; tomato spotted wilt virus, which re-emerged affecting more than 800 plant species; yellow mottle virus disease, affecting irrigated rice^4^; and recently, the cassava mosaic geminiviruses that are threatening cassava production^5^. The emergence and spread of infectious viral diseases is driven by intrinsic viral and host factors, in addition to ecological, agronomical and socioeconomical factors^6–9^. Climate change is likely to increase the frequency of emerging viral diseases in plant crops^10–12^. Warming and highly variable climate may directly and indirectly affect host, vector, and viral traits, and further influence viral epidemics both in cultivated and wild plants. Global climatic models predict a substantial rise in temperature with unusual temperature variations. This could change the distribution and behavior of viral vectors and/or provide optimal climatic conditions for pre-existing or novel viruses in new geographical areas^13^. Environmental heterogeneity, which has been recently shown to shape plant-virus infection networks at temporal and spatial scales seems to be a key driver of the disease emergence and dynamics^14^. Therefore, understanding how and, to what extent, temperature variations affect the eco-evolutionary dynamics of viral populations is essential. This will help us to understand the effect of global warming on virus emergence and epidemiology.

Multiple viral infections (i.e. mixed infections) are gaining considerable significance on plant crops. Hence, within-plant virus-virus interactions can have causal effects on epidemiology as a result in competitive interactions during an infection^15–19^. For example, competitive interactions can have major consequences on virulence and fitness of the viruses^19–21^. The viral interactions may also create a spatiotemporal distribution pattern of viral populations through the host, and thus, structured sub-populations can coexist^6,22–24^. The competition for common host resources is likely to affect the accumulation and spread of each virus allowing a spatial structuring of the viral population. This forms a complex genotype mosaic through the host that may further display asynchronous variation in competition through time^25^. This spatial structure of populations has been theoretically predicted to affect mutation rates and increase diversifying selection through a competition/colonization trade-off^22,26–28^. Thus, depending on the mutation rate and effective population size^29^, each viral population will have more or less ecological success to explore the fitness landscape^30^. This may leave a genetic signature for at least one of the viruses involved in the infection. The effect of mixed infections on the population genetic variability could be largely contingent upon host ecology and environmental changes. However, despite that this is a epidemiologically relevant situation, it is still a neglected topic. The potential effect of the within-host interactions between strains / genotypes of the same virus in changing environmental conditions has yet to be proven.

Here, we used two strains of pepino mosaic virus (PepMV; genus *Potexvirus*, family *Alphaflexiviridae*); the European (EU) and Chilean (CH2)^31,32^. PepMV causes one of the most important viral diseases in tomato worldwide, producing subsequent outbreaks over recent years^33^. PepMV genome consists of a positive-sense, single-stranded RNA molecule of approximately 6.4 kb in length, containing five open reading frames (ORFs) flanked by two untranslated regions (UTRs), with a poly(A) tail at the 3’ end of the genomic RNA (gRNA). PepMV diversity is mostly composed by isolates of EU and CH2 strains. CH2 strain has become the dominant type, but EU strain still occurs mainly in mixed infections in greenhouse tomato crops^31,32,34^.

Our aim was to understand the interplay between mixed viral infections and a fundamental niche dimension, temperature, in order to understand the effect of climate change on viral disease emergence. We hypothesized that growth temperature affects virus-virus interactions, which in turn shape the within-host genetic diversity of viral populations as a consequence of a competition/replication trade-off. To test this, we experimentally combined the effect of the temperature with the potential effect of virus-virus interactions *in planta.* In addition, we introduced mathematical models that incorporated temperature-dependent replication rates to understand (*i*) the interplay between temperature and viral replication and the strength of competition in mixed infections, and (*ii*) provide a mechanistic model able to qualitatively reproduce the observed dynamics and make further predictions.

## RESULTS

### Identifying significant drivers of PepMV accumulation

To investigate how growth temperature (*T*), type of infection (*I*) and genetic differences among PepMV isolates (*A*) affected gRNA accumulation in tomato plants 7 and 60 dpi (Fig. 1A), we fitted the viral load (*VL*) data to Equation (5). Table 1 shows the statistical summary of the corresponding GLM analysis. Only the interaction term *A*×*T* was not significant. However, a number of terms in Equation (5) (highlighted in gray in Table 1) had low statistical power (1 − *β* < 0.800), thus wrongly failing to reject the null hypothesis (type II error), and the magnitude of the effect classified as small according to the 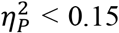 criterion. Thus, hereafter we comment on highly significant and relevant terms. Firstly, regarding the main factors, only the *A* and *T* were relevant by themselves to explain the observed variability in *VL* (Table 1, Fig. 1B and 1C). Overall, CH2 accumulates to higher concentrations than EU, with an averaged accumulation smaller at 30 °C. The covariable duration of infection (*t*) also had a net effect on *VL*, supporting that it increased between 7 and 60 dpi (data points in Fig. 2 and Fig. 3). Remarkably, the interaction between the three main factors and *t* was also highly significant, indicating that the observed effects changed with time. While *VL* was 3.1-fold larger for EU 7 dpi, it was 8.0-fold larger for CH2 60 dpi. *VL* was, on average, larger in single than in mixed infections, but the magnitude of this difference slightly decreased along the progress of infection (1.8-fold at 7 dpi, and 1.3-fold at 60 dpi). At 7 dpi, infections at 20 °C were 3.3 times more productive than a 30 °C, slightly less (2.7-fold) at 60 dpi.

**Figure 1.**
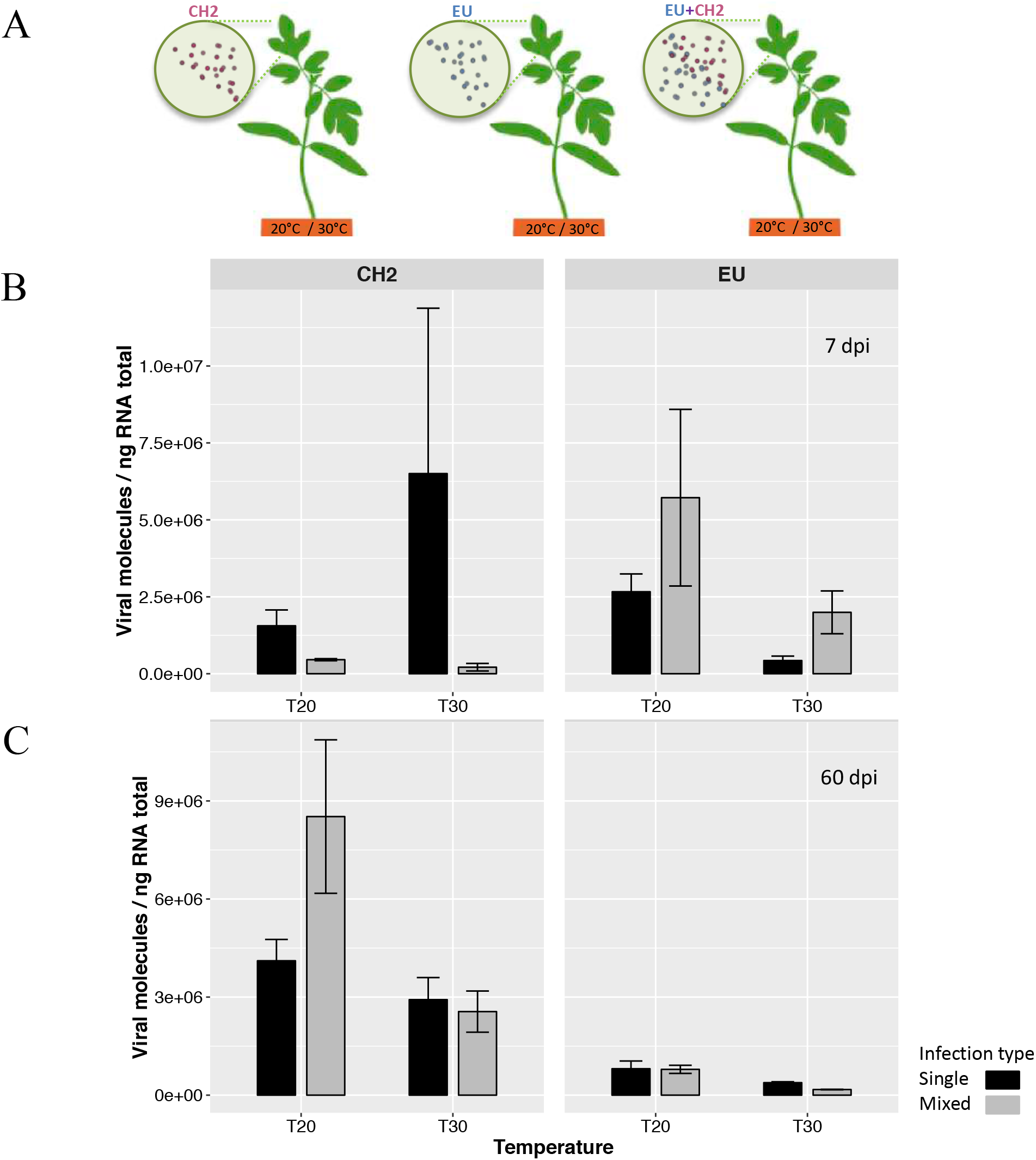
(**A)** What is the effect of mixed viral infections on the fitness and genetic diversity of viral populations within a host? **(B and C)** Barplots exhibiting the viral load (RNA molecules / ng RNA total) of each PepMV (CH2 and EU) strain in tomato plants grown at 20 °C and 30 °C under single (black) and mixed infection (grey) condition. Viral accumulation was inferred by absolute quantification using RT-qPCR after 7 dpi (**B**) and 60 dpi (**C**).

**Figure 2.**
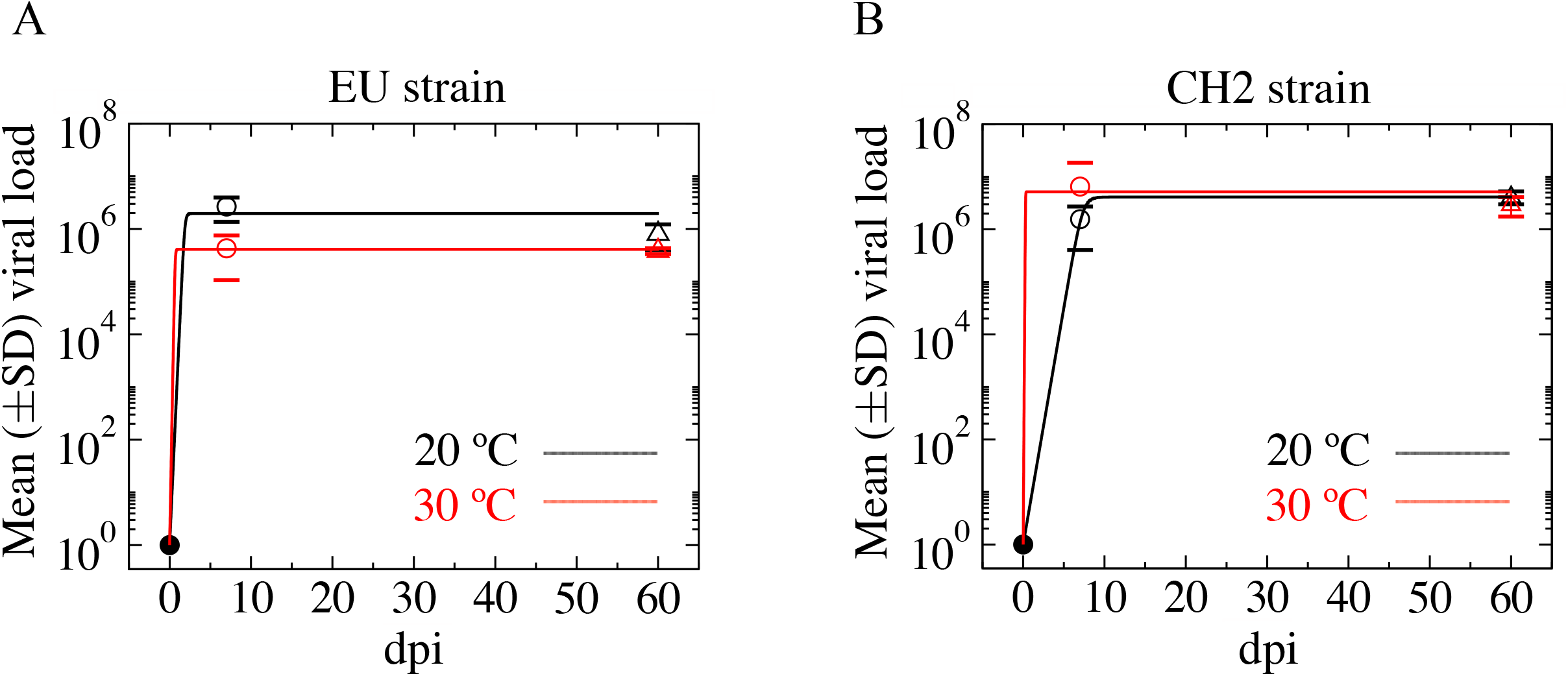
Fitting of the mathematical model (solid lines) to the experimental data (circles) for single infections for strains EU (**A**) and CH2 (**B**) at 20 °C (black) and 30 °C (red). Panels display the fit to the mean values (± SD) of the experimental data. The same fitting over the replicates is shown in Fig. S5. The dynamics of the model have been obtained using the best parameter set found by the Macroevolutionary Algorithm (see Section 3 and Fig. S6 in the SI for the same analyses using the mean optimized parameters and mean best parameter vectors).

**Figure 3.**
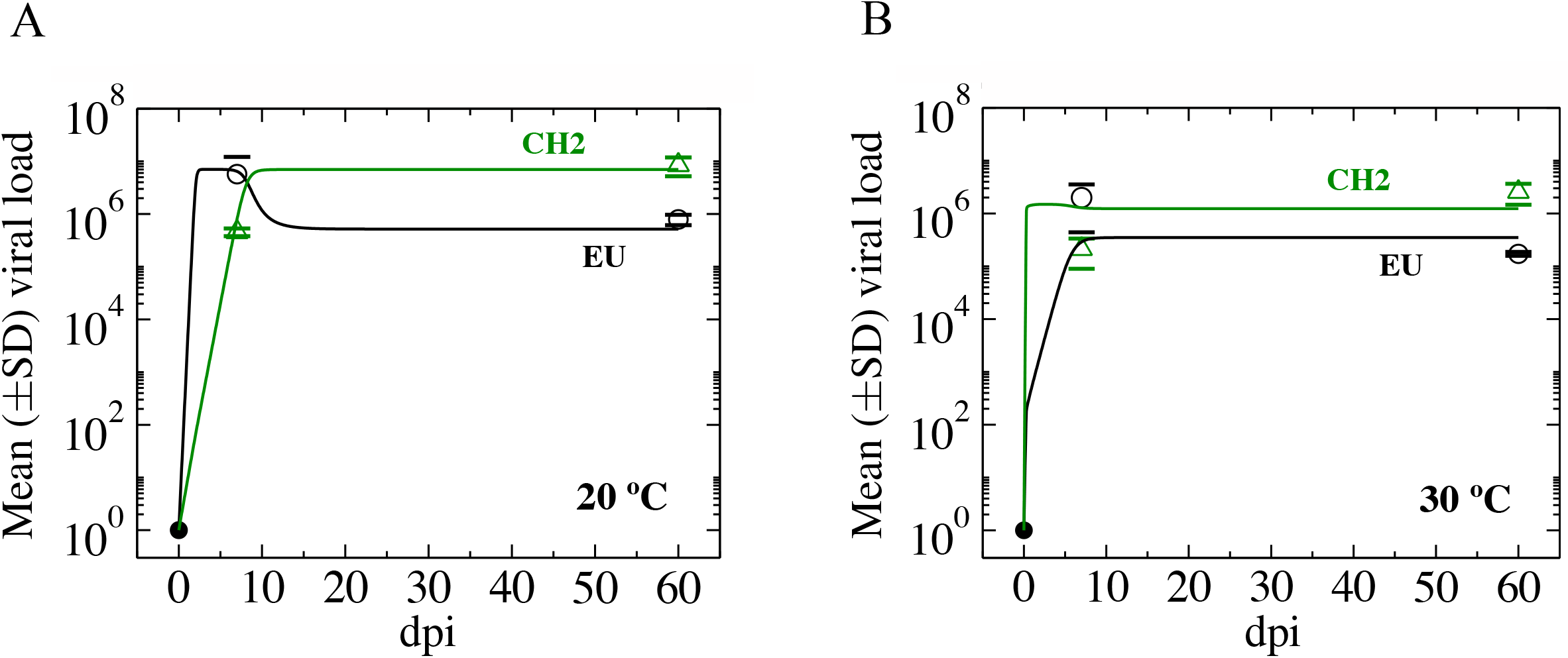
Fitting of the mathematical model with competition between viral strains to the experimental data for mixed infections at 20 °C (A) and 30 °C (B) using the parameters providing the best bit of the Macroevolutionary Algorithm. Panels display the fit to mean (± SD) experimental values. Experimental data is shown with circles while the time series obtained with the model are displayed with solid lines. Black and green data correspond to data for EU and CH2, respectively. The same fitting over the replicates is displayed in Fig. S9 (see Section S4 and Fig. S10 for the same analyses using the mean optimized parameters and mean best parameter vectors).

**Table 1.** Results of the GLM fitting of the viral load data to Equation (5).

| Source of variation | LRT<sup>1</sup> | df | *P* | $\eta_p^2$ | $1 - \beta^2$ |
| --- | --- | --- | --- | --- | --- |
| $\lambda$ | 13919360.545 | 1 | < 0.001 | 0.993 | 1.000 |
| *t* | 128.340 | 1 | < 0.001 | 0.483 | 1.000 |
| *A* | 298.137 | 1 | < 0.001 | 0.455 | 0.889 |
| *t*×*A* | 417.126 | 1 | < 0.001 | 0.913 | 1.000 |
| *I* | 129.466 | 1 | < 0.001 | 0.165 | 0.342 |
| *t*×*I* | 67.675 | 1 | < 0.001 | 0.352 | 1.000 |
| *T* | 294.315 | 1 | < 0.001 | 0.410 | 0.826 |
| *t*×*T* | 70.774 | 1 | < 0.001 | 0.352 | 1.000 |
| *A*×*I* | 280.915 | 1 | < 0.001 | 0.416 | 0.836 |
| *t*×*A*×*I* | 260.109 | 1 | < 0.001 | 0.750 | 1.000 |
| *A*×*T* | 3.630 | 1 | 0.057 | 0.003 | 0.054 |
| *t*×*A*×*T* | 12.626 | 1 | < 0.001 | 0.044 | 0.649 |
| *I*×*T* | 71.393 | 1 | < 0.001 | 0.071 | 0.161 |
| *t*×*I*×*T* | 8.810 | 1 | 0.003 | 0.025 | 0.422 |
| *A*×*I*×*T* | 146.124 | 1 | < 0.001 | 0.176 | 0.363 |
| *t*×*A*×*I*×*T* | 81.264 | 1 | < 0.001 | 0.391 | 1.000 |
| *R*(*A*×*I*×*T*) | 511.786 | 14 | < 0.001 | 0.857 | 1.000 |
| *t*×*R*(*A*×*I*×*T*) | 257.093 | 14 | < 0.001 | 0.779 | 1.000 |

<sup>1</sup>Likelihood ratio test, asymptotically distributed as a $\chi^2$ .<sup>2</sup>Power of the test. Shadowed terms are not further considered because their low statistical power ( $1 - \beta < 0.800$ ) or small magnitude effect ( $\eta_p^2 < 0.150$ ).

Secondly, the only relevant pairwise interaction was *A*×*I* (Table 1, Fig. 1B and 1C): EU accumulated 1.6-fold more in mixed than in single infections, whereas for CH2 the situation was the reverse: it accumulated 4.2-fold more in single than in mixed infections. Indeed, these differences strongly depend on *t*: while EU productivity in mixed infections was 2.6-fold higher 7 dpi, it was only 1.9-fold higher 60 dpi; in the case of CH2, it accumulated 8.9-fold more in single infections 7 dpi but 1.2-times more in mixed infections 60 dpi.

Thirdly, the tree-ways interaction between the main factors was not significant *per se,* but only throughout a dependence with the duration of infection (Table 1 and data points in Fig. 2 and Fig. 3). In the case of EU, *VL* at 7 dpi was larger for mixed infections (the precise magnitude depending on *T*), while at 60 dpi it was larger for single infections (again, the precise magnitude depending on *T*). The opposite pattern was observed for CH2: *VL* was larger in single infections 7 dpi and slightly larger in mixed infections after 60 dpi at 20 °C, yet the reverse was found at 30 °C.

### Dynamical modelling and parameters estimation for single infections

The mathematical models we have used, albeit simple, reproduced the full dynamics of the experiments considering the data at 7 and 60 dpi as part of the same time series. Also, despite lacking a detailed description of the virus replication cycle and spread over the plant, the models also provided quantitative information on key virological parameters for the different experimental conditions. As mentioned, the dynamics of the model for single infections is summarized in Section S1.1 in the SI. The estimated replication rates (computed from the estimation of parameters *a* and *b* in Equation (2)) providing the best fits for the single infections with the EU strain were *r_E_*(20) = 7.760 ng gRNA/day and *r_E_*(30) = 19.012 ng gRNA/day. The carrying capacity, *K*, for the EU strain was found higher at the lower temperature [*K*(20) = 1.968 mg gRNA], compared to the experiments at higher temperature [*K*(30) = 0.412 mg gRNA]. Therefore, by increasing *T*, EU replication rate increases 2.45 times, while the total amount of gRNA produced is 4.78 times lower, in agreement with data shown Fig. 1B. The fitting of the mathematical model to the experimental data (mean values) for single infections at both temperature conditions are shown in Fig. 2A. Notice that the initial replication phase is extremely fast, with the equilibrium value being achieved even before 7 dpi. Similar fitting were observed in the experimental replicates Fig. S5a, in addition to the fittings with the mean values of the optimised and best perameters among 50 replicates of the MA (Fig. S6a). After longer times of infection, the populations of EU and CH2 remain approximately at the same *VL* (Fig. 2).

Concerning CH2 strain, estimated replication rates with better fits were: *r_C_*(20) = 2.076 ng gRNA/day and *r_C_*(30) = 53.923 ng gRNA/day. Here, the carrying capacities remained at similar ranges, with *K*(20) = 4.110 mg gRNA and *K*(30) = 5.160 mg gRNA. The fit of the model with these parameters is displayed in Fig. 2B, and similar fitting was obtained using the mean experimental values (Fig. S5B). Here, the dynamics was similar to the one of EU strain. Equilibrium was achieved extremely fast and then the *VL*s remained quite stable until 60 dpi. By increasing *T*, CH2 replication rate increased 25.97 times, while the total amount of gRNA produced remained on the same order of magnitude at both temperatures (1.26-fold increase), in agreement with *VL* data shown in Fig. 1B and 1C. The fitting performed with the mean values of the optimised parameters and of the best parameters sets is displayed in Fig. S6b.

Finally, the functional relationship described by Equation (2) allowed us to obtain the thermal reaction norm for each strain from the estimation of parameters *a* and *b* for single infections. The results, displayed in Fig. S8 for the best fits and the mean estimations obtained after the MA optimization, indicated that both EU and CH2 shared qualitatively similar profiles, with a linear increase in replication rate by temperature and a marked decline after 45 °C, see Section 3.5 of the SI for further details. These results indicate that *T* had a positive effect in the replication rates of both PepMV types, although the effect was much larger for CH2 than for EU. The effect on maximum *VL* (measured as *K*) went in different directions with *T*: decreases for EU but increased for CH2.

### Dynamical modelling and parameters estimation for mixed infections

To investigate the impact of competition between both strains at the levels of population dynamics and temperature we used Equations (3) and (4). The dynamical behaviour of the model with competition can be found in Section 1.3 in the SI. Following the same procedure than for single infections, we fitted the mathematical model with competition to the experimental data also using the MA optimization. In order to quantify the competition coefficients *β_EC_* and *β_CE_* we used the replication rates obtained with the best fits for each strain at different temperatures in the single infections. By doing so, and given the structure of the mathematical model, we assumed that changes in the replication of both strains, and thus in accumulated *VL*, were given only by competitive effects. Also, we estimated again *K* for two reasons: (*i*) the variability of this parameter obtained in the single infections experiments at different temperatures did not allow us to choose a proper (constant) value and (*ii*) an antagonistic interaction between the two strains during mixed infections could negatively affect total viral accumulation^16^. The mathematical model with competition provided the best fit of the experiments at 20 °C for parameters *β_EC_* = 0.932, *β_CE_* = 0.079 and *K* = 7.042 mg gRNA. We showed that both viral types were able to coexist at 20 °C despite CH2 competed better than EU (*β_EC_*/*β_CE_* = 11.80 times stronger). The results of the fitting with these parameter values to the mean experimental values are displayed in Fig. 3A (the same fitting on the replicates is provided in Fig. S9, while the fitting of the model with the mean values of the optimised and best parameters is displayed in Fig. S10a). We note that at low *VL* EU dominated over CH2. However, once larger population numbers are achieved, CH2 continues accumulating while EU substantially decreased.

Similar results were found at 30 °C (fitting with the best parameters is shown in Fig. 3B (mean values), in Fig. S9b on the replicates, and in Fig. S10 performed with the mean values of the optimised and best parameters). Under this temperature, the parameters giving the best fit were *β_EC_* = 0.929, *β_CE_* = 0.750 and *K* = 1.497 mg gRNA. Hence, at 30 °C we found that coexistence at this higher temperature was also possible and, more remarkably, now CH2 would interfer with EU less efficiently (*β_EC_*/*β_CE_* = 1.24) than at 20 °C.

### Mutation occurrence from full-length genome sequencing

To examine the influence of differences in replication and competition between CH2 and EU on the viral genetic variability within-populations, we first identified a genomic region at the 3’ end of the viral gRNA that allows distinguishing both PepMV types and full-length genome amplifications from mixed infected samples. This provided us to obtain whole-genome sequences of each PepMV populations after 60 dpi, as well as we included the inocula. After cleaning and trimming the raw NGS data we obtained between 809,756 and 1,777,280 reads per sample, resulting in a theoretical average coverage ranging from 31,750× to 69,687×. These reads were further filtered out by quality, and using the initial inocula as reference, we identified SNPs (and evaluated their frequency in the samples) that appear during the infection experiments. After, excluding SNPs present in the inocula and considering variants with a frequency higher than 0.01, including those variants found in the overlapping region between TGB2 and TGB3 genes as double variants because of the potential effect in both proteins, we end up with a total of 519 SNPs (Table S1, Fig. 4A and Fig. 4B). No indels or recombination events were observed. Attending to the frequency of the new mutations, the vast majority were ranging from 0.01 to 0.1, with only 3.28% of the total SNPs displaying frequencies > 0.5 (Table S1). Note that all these high-frequency mutations were specific to the CH2 populations.

**Figure 4.**
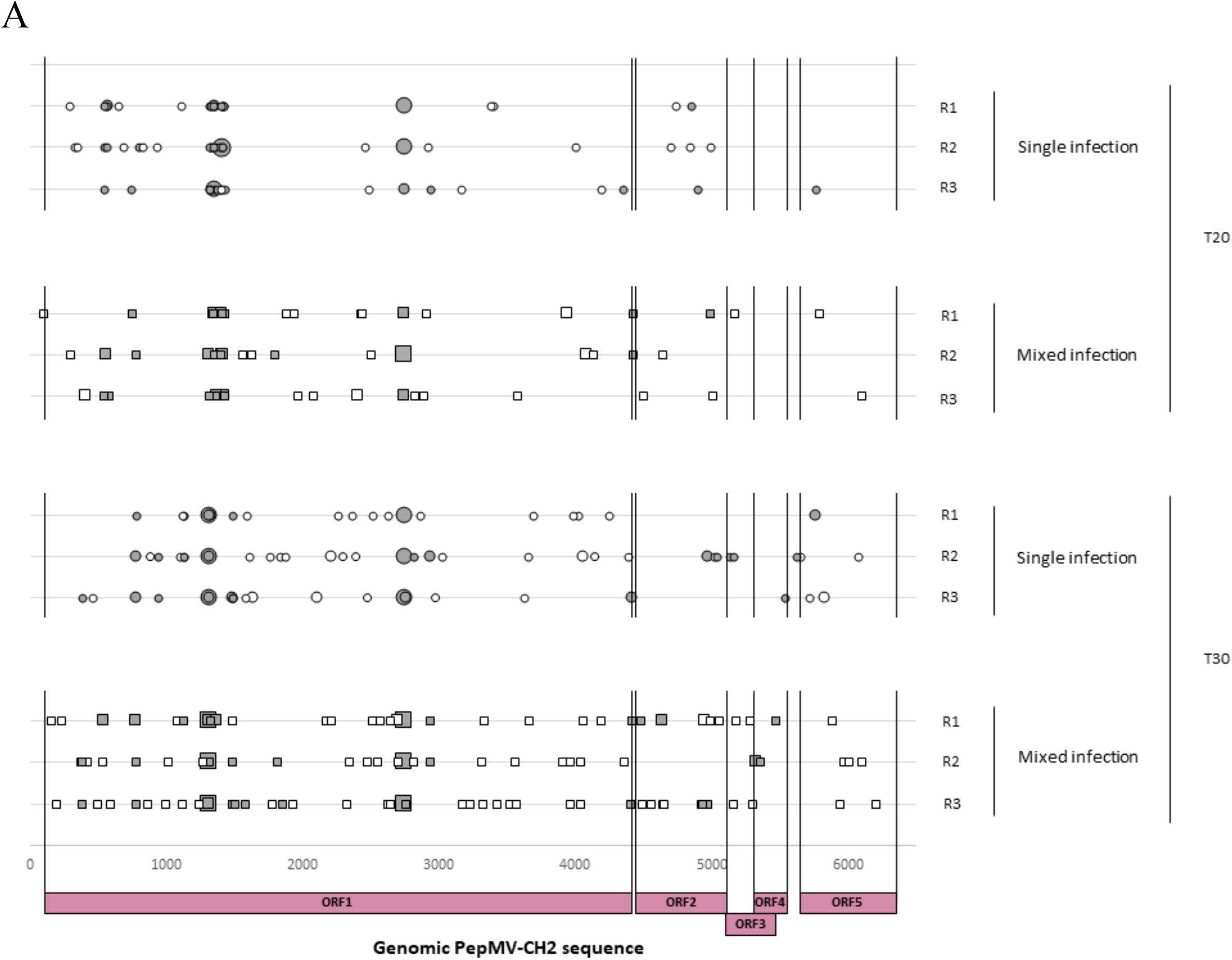
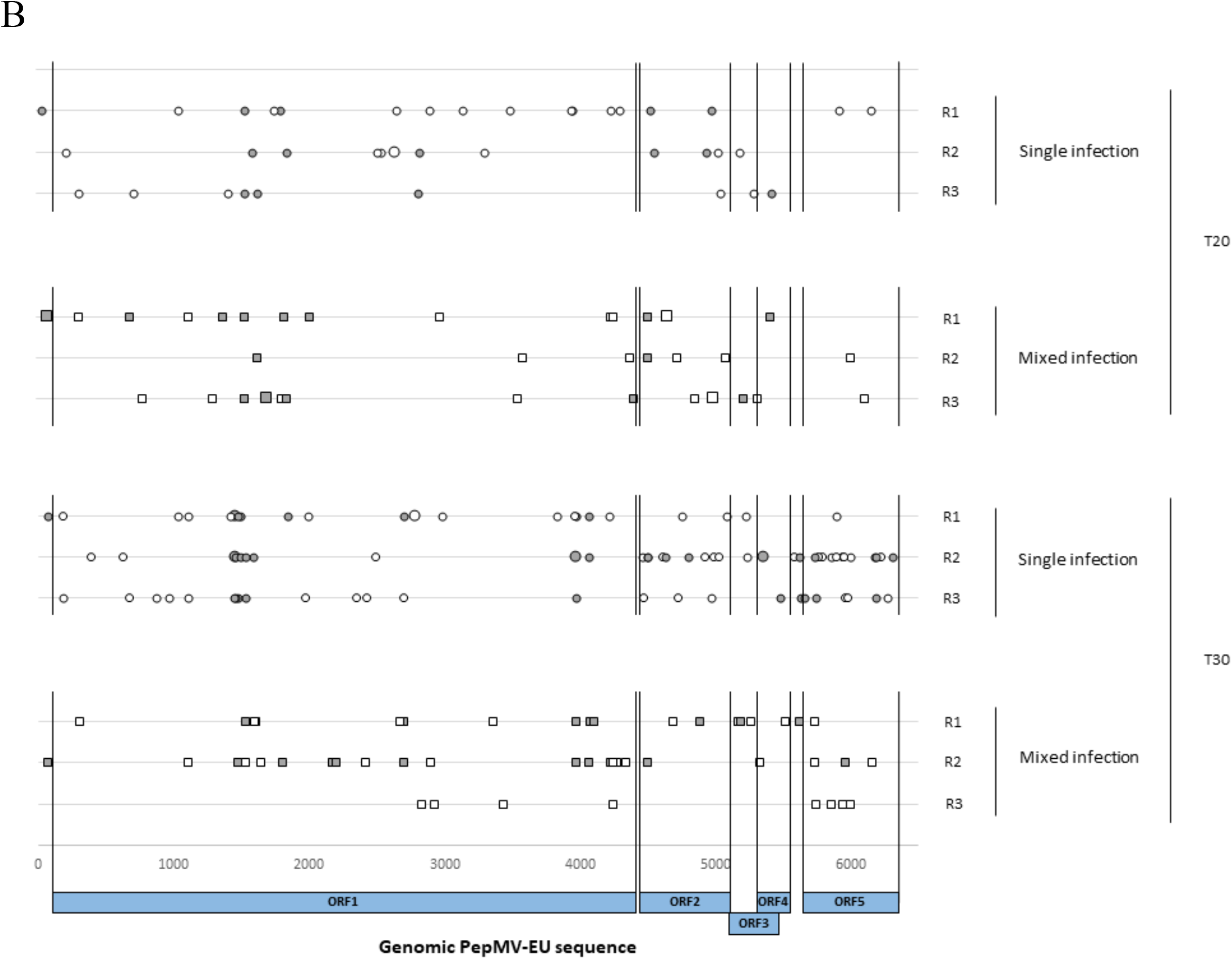
Occurrence and distribution of sites exhibiting mutations in the full-length genome of both PepMV isolate populations; (**A**) PepMV-PS5 (CH2 type) and (**B**) PepMV-Sp13 (EU type). The SNPs are marked (○; synonymous mutations) and (**●**; non-synonymous mutations) for each replicate (n=3) in single and mixed infections under 20 °C and 30 °C conditions.

SNP counts (*SC*) of PepMV within-host populations are summarized in Fig. 5. These data were fitted to the model described by Equation (6) and the results of the analyses are shown in Table 2. Overall, and focusing first on the main effects, highly significant differences of large magnitude existed between the average number of mutations fixed by the CH2 and EU within-host populations. CH2 populations accumulated 1.57-fold higher mutations than the EU populations (Fig. 5). No differences existed among single and mixed infections (*I* term in Table 2) in terms of the number of mutations accumulated. The average number of mutations per within-host population also significantly increased 1.57-fold with temperature (*T* term in Table 2 and Fig. 5). The higher-order terms *A*×*I* and *A*×*I*×*T* in Equation (6) were also significant (*P* ≤ 0.011) and their effects could be considered of large magnitude 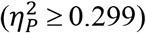, suggesting that the observed effect associated with the viral genotype indeed would depend on its interaction with the type of infection and temperature in a nonlinear way. However, given that the power of the two tests was 1 − *β* < 0.8, this conclusion should be taken carefully (Table 2). Furthermore, genetic diversity of CH2 and EU populations was evaluated by using heterozygosity (*H*) and nucleotide diversity (*π*) estimators (Table 3). We found that both, *H* and *π*, were higher in the CH2 than EU populations, with averaged *H* values higher at 20 °C in mixed infections. However, averaged *π* values were higher at 30 °C in mixed infections of the CH2 populations that, in turn, was consistent with significant increases of the numbers of synonynous than non-synonymous mutations (Table 3). Hence, conservatively, we can conclude that populations of both PepMV types differed in their mutational load in an infection- and temperature-dependent manner.

**Figure 5.**
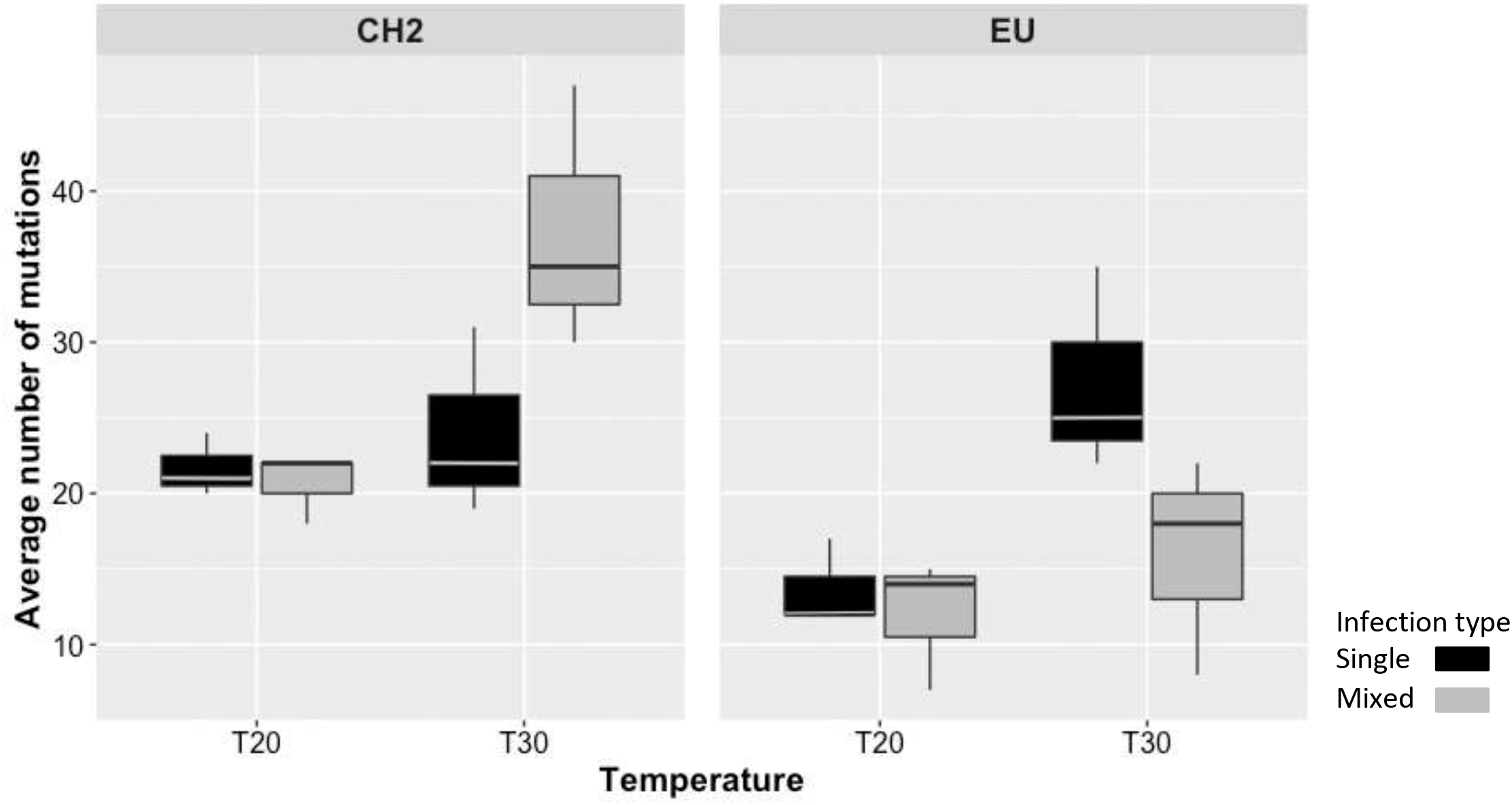
Boxplot displaying the average number of SNPs for each PepMV-CH2 and - EU population in single (black) and mixed infections (grey) at 20 °C and 30 °C of plant growth temperature. The horizontal line shows the median value, and whiskers show the minimum and maximum values.

**Table 2.** Results of the GLM fitting of the mutation count data to Equation (6).

| Source of variation | LRT<sup>1</sup> | df | *P* | $\eta_p^2$ | $1 - \beta^2$ |
| --- | --- | --- | --- | --- | --- |
| $\sigma$ | 1416.585 | 1 | < 0.001 | 0.955 | 1.000 |
| *A* | 23.662 | 1 | < 0.001 | 0.478 | 0.949 |
| *I* | 0.403 | 1 | 0.526 | 0.000 | 0.050 |
| *T* | 23.155 | 1 | < 0.001 | 0.514 | 0.971 |
| *A*×*I* | 6.993 | 1 | 0.008 | 0.299 | 0.690 |
| *A*×*T* | 0.991 | 1 | 0.320 | 0.000 | 0.050 |
| *I*×*T* | 0.019 | 1 | 0.890 | 0.013 | 0.072 |
| *A*×*I*×*T* | 6.457 | 1 | 0.011 | 0.311 | 0.713 |

<sup>1</sup>Likelihood ratio test, asymptotically distributed as a $\chi^2$ .<sup>2</sup>Power of the test. Shadowed terms are not further considered because their low statistical power ( $1 - \beta < 0.800$ ) or small magnitude effect ( $\eta_p^2 < 0.150$ ).

**Table 3.** Average number of SNPs and estimation of genetic variability indexes (*π*, nucleotide diversity and *H*, observed heterozygosity) for each viral strain population grouped by the temperature and type of infection. Mutations are classified according to the type of change [synonymous (syn) and non-synonymous (nsyn), as well as non-coding variants (nc)].

| PepMV strain | Temperature (°C) | Type of infection | syn | nsyn | nc | Total | $\pi (\times 10^{-4})$ | SD ($\pi$) ($\times 10^{-4}$) | $H$ | SD ($H$) |
| --- | --- | --- | --- | --- | --- | --- | --- | --- | --- | --- |
| CH2 | 20 | Single | 21 | 44 | 0 | 65 | 4.584 | 0.240 | 0.090 | 0.008 |
|  |  | Mixed | 26 | 33 | 2 | 61 | 5.400 | 0.408 | 0.107 | 0.016 |
|  | 30 | Single | 37 | 33 | 2 | 72 | 6.591 | 2.399 | 0.079 | 0.043 |
|  |  | Mixed | 71 | 35 | 8 | 114 | 6.910 | 1.335 | 0.057 | 0.023 |
| EU | 20 | Single | 25 | 13 | 1 | 39 | 3.549 | 0.758 | 0.049 | 0.019 |
|  |  | Mixed | 20 | 15 | 1 | 36 | 4.264 | 0.555 | 0.071 | 0.018 |
|  | 30 | Single | 50 | 31 | 3 | 84 | 5.101 | 1.191 | 0.065 | 0.013 |
|  |  | Mixed | 29 | 17 | 2 | 48 | 3.345 | 0.562 | 0.043 | 0.005 |

## DISCUSSION

Ongoing global warming is expected to increase the incidence of plant pests^12,35,36^. At the same time, accumulating evidence shows that plant viral diseases frequently contain mixed strain infections with a high prevalence in plant crops. Our interest was focused on the effect of growth temperature rise in combination with mixed viral infections. In this study, we connected both abiotic and biotic factors, and more explicitly, sought to explore the relationship between temperature and evolutionary dynamics of viral populations in single and mixed infections. To do so, we combined the experimental approach to mathematical modelling. PepMV-CH2 type experienced a fitness cost in the presence of EU type during the mixed experimental infections in tomato plants, which is consistent with previous studies^31,34^. In these previous studies, the asymmetrical antagonistic interaction between these two strains provided a partial explanation for the maintenance of the genetic structure of the PepMV populations in tomato. Both strains have been co-occurring in mixed infections and contributed to the evolutionary dynamics of PepMV populations^31^. However, sequencing data from several field isolates hardly inform about the type of virus-virus interactions that may be modulating PepMV genetic variation in plants. Agricultural practices and environmental factors may also affect virus’ genetic diversity. In our controlled experiment, we found that the magnitude and sign of the interactions between the viral strains appeared to vary during the infection, and substantially changed when tomato plants grew at 30 °C, compared to the 20 °C temperature conditions (Fig. 1). This proved the first part of our hypothesis. We suggest that the increasing growth temperature alters host plant physiological and biological traits that in turn allow changes in virus multiplication. Therefore, increasing temperature may influence the accumulation of individual viruses co-infecting the same host plant and affect their potential viral interactions.

We found that the strength of viral competition was relaxed in a temperature-dependent manner, which had an effect on the nucleotide variation of both of the viral strains, depending on the type of infection (Fig. 5). The same relaxation was identified and quantified with the mathematical modelling as well. Therefore, we can accept the second part of our hypothesis. Compared to the single infection, we observed that the genetic variation of CH2 population increased in the mixed infections at 30 °C while the opposite pattern was found for EU population (Fig. 5). Interestingly, this increase in the genetic variation of CH2 seems to be related to a weaker interference with EU at 30 °C (Fig. 5 and Fig. S8), and also to higher replication rates for CH2 and possibly a higher mutational load as well. Thus, increasing temperature could modify viruses’ evolution rates in mixed infections. Higher virus multiplication and reduced selection pressure seem to lead to an increase in the viral genetic variability of within-host populations. This dynamics seem to be similar to that observed in crop production studies during seasonal heat events; heat has been associated to an increase in viral multiplication, systemic movement and transmission^10,11,37^. As a summary, temperature likely contributes to the origin and maintenance of the viruses’ genetic diversity by the remission of viruses’ competitive interactions in mixed infections. In this sense, host plant’s phenotypic plasticity and resilience to environmental changes can be playing important roles in how climate change will affect virus-host interactions. These changes are likely to modulate the viral genetic diversity in a continuum between single and mixed infections. This suggest that our framework should be extended on how host plants’ tolerance to warmer temperatures affect the viral genetic variability and competitive interactions in mixed infections. Further research could help in anticipating the impact of plant breeding programs for heat tolerance and how new plant cultivars will drive the eco-evolutionary dynamics of viral populations in plant crops.

Most of the information on virus evolution relies on the use of single infections. They allow qualitative and, to a much lesser extent, quantitative interpretations about the emerging viral diseases, as mixed viral infections are common in nature. Viral occurrence and prevalence in plant crops seems to be largely explained by either neutral, synergistic or antagonistic interactions among co-infecting viral species^15–18^. However, these viral interactions do not recognize the potential genetic plasticity to strain-interactions co-infecting the same plant. It is possibly because of the limitations of viral systems that allow address whole-genome sequencing analysis, as different strains or the same virus appear to have larger nucleotide similarity. This particular bias happened with both CH2 and EU strains. It was elucidated by using a genomic region at the 3′ end of the viral RNA genome that allowed to obtain the full-length genomes from mixed infections, capturing the nucleotide temporal variation of highly related plant viruses. First, we report that while mixed infections of PepMV genotypes may set a recombination occurrence, no recombinant variants were found in the present experimental assay, neither in the previous field tomato surveys since 2005^31^. This suggests that recombinants appeared to be not selected in the virus populations. Second, our mutational analysis showed that SNPs was the only mechanism contributing to the genetic variability within PepMV populations. Third, we found that the type of infection (single and mixed) did not influence the average population mutational load at 20 °C. The last observation is in accordance with previous studies where the viral population diversity has not seemed to differ between mixed and single infections^38,39^. However, the higher temperature appeared to change the situation. Our results revealed that both strains displayed higher genetic variation and greater differences in accumulation at higher temperatures (Fig. 4 and 5). In particular, the CH2 population had higher number of SNPs in mixed than in single infections at higher temperature. The increased effective population size in mixed infections at 30 °C could in part explain this strain-specific mutational effect.

We observed that CH2 and EU replication rates increased by increasing temperature, with a replicative effect greater on CH2. Other theoretical studies have shown accordingly the importance of competitive ability and viral fitness trade-off in determining the evolutionary dynamics of the viral populations^26,40^. We speculate that a trade-off between the replication rate and competition has also influenced this different mutational effect. Under the same framework, it has been theoretically demonstrated that the trade-off between replication and competition can affect diversity, with moderate trade-off promoting diversity^41^. On one hand, a moderate trade-off could have boosted the viral genetic diversity observed in this study, as replication rate of the CH2 was higher than EU at 30° C, without any increase of the viral load, which suggests a reduced competitive ability. On the other hand, it is likely that EU population had an increased cost between replication and competition in mixed infections under high temperature reducing its genetic variability compared to single infections. Different viral combinations with different fitness responses (see examples in ref. 18) may modify the outcomes of this eco-evolutionary dynamics, thus emphasizing the need of further research.

What is the extent of the biotic selective pressure on viral genetic diversity? It has been suggested that the viral mutational load would be purged faster in a mixed infection than in a single infection due to reassortment and selection^42^. However, other studies have showed the opposite, as deleterious mutants were purged slower in mixed than single infections showing less robustness in mixed infections^39,42^. Once new variants emerge in a population, they could be separated into different cell tissues and result a within-host evolution through a mutation-selection balance^42–45^, with their potential maintenance by negative-frequency-dependent selection^41^. In case a colonizer is very rapidly colonizing the available cells and the competitor has a greater survival capacity in co-infected cells, a co-existence can be expected. Morevoer, the multiplicity of infection of each virus could also affect the spatiotemporal distribution of multiple viruses co-infecting the same plant, and hence, influencing viral diversity and their adaptive evolution at the cellular level^23,26,46–49^. Therefore, genetic trade-off between colonizer and competitor in a viral infection could lead to an additional evolutionary advantage of rare variants, and selection may favor genotypes that require fewer stabilizing effects to allow coexistence^50^. All these aspects emphasize the lack of a comprehensive understanding of the genetic and ecological bases of the evolutionary dynamics of viral populations in mixed infections.

In conclusion, the observed tight connection between the temperature and viral interactions in mixed infection leads to the expectation that future warming will result in high viral diversity in plants. This genetic diversity may be modulated by changing within-host viral interactions. Due to the changing virus-virus interactions, the rising temperature could impose distinct abiotic selection pressures on viral populations in mixed infections. This could affect the accumulation and replication rate of viruses within a host, which could, in turn, affect the viral genetic population structures and shape their evolutionary dynamics. Our results can readily be applied to other pathogen systems caused by mixed infections. A noteworthy remark is that pathogens evolve not only within hosts but also between hosts, and the evolution seems to depend on the frequency of mixed infections and the success of re-infections in one host. Assessing the mechanistic basis of selective differences will allow us to understand the importance of abiotic and biotic factors in wider pathosystems.

## MATERIALS & METHODS

### Plant growth conditions, virus inoculation and sampling

Two infectious PepMV clones belonging to the PepMV-Sp13 isolate (EU type)^51^ or PepMV-PS5 isolate (CH2 type)^34,52^ were used to agroinfiltrated *Nicotiana benthamiana* Domin plants. After 14 days post-inoculation (dpi), viral particles were purified from the homogenized plant tissue, following a series of centrifuges and a PEG precipitation^53,54^. Then, tomato plants (*Solanum lycopersicum* L. cv. Money Maker) were grown in a greenhouse with a photoperiod of 16 h light:8 h dark and a temperature between 22-26 °C with a day/night cycle. After 30 days post-germination, two sets of 33 plants were placed into a greenhouse with temperature conditions of 20 °C or 30 °C. For each temperature condition, six tomato plants were mock-inoculated and nine plants per treatment were mechanically inoculated by PepMV-Sp13 (EU type), PepMV-PS5 (CH2 type) or mixed infections of both isolates. The inoculations were performed on the third and fourth true leaves by rubbing Carborundum and a suspension of virions particles at 500 ng/μL in sodium phosphate buffer (30 mM). During the experiment, PepMV infection was checked by the detection of both PepMV types from a leaf-sample of each plant collected at 12, 30 and 58 dpi. Total RNA was extracted using Tri-reagent, and 1μL of RNA was placed on a nylon membrane and fixed with ultraviolet light (CL-1000 Ultraviolet Crosslinker). PepMV infections were detected by dot-blot molecular hybridization using specific RNA probes to detect EU and CH2^55^ and following the Roche protocol (RNA labelling and detection kit). Finally, the membrane was revealed using a chemiluminescent detector Amersham™ Imager 600.

### Full-length viral amplification

Plant material was collected 60 dpi. The last eight apical-levels of plants were harvested and grinded in a mortar using liquid nitrogen. Three pools were made per treatment, with three plants per pool. A sample from each pool was taken and RNA extraction was performed using RNeasy Plant Mini Kit (QIAGEN) and stored at −20 °C until use. One thousand ng of total RNA was used for retro-transcription (RT) (Expand™ Reverse Transcriptase, Roche). In the case of mixed infections, RT reactions were performed in duplicate, one of them using a specific primer for the EU strain (EU_rv: 5’-TTTTTTTTTTTTTTATTTCAAAGAAATAATTA-3’) and another using a specific primer for the CH2 strain (CH2_rv: 5’-TTTTTTTTTTTTTAGTAGATTTAGATACTAAG-3’). Then, cDNA was amplified using Phusion High-Fidelity DNA Polymerase (Thermo Fisher Scientific) and specific primers for each isolate (EU_rv and EU_fw: 5’-CGCGGATCCGGAAAACAAAATAAATA AATAAATATAC-3’ for EU and CH2_rv and CH2_fw: 5’-GAAAACAAAACATAACACATAATATCAAAAGTGACC-3’ for CH2). The initial denaturation was done at 98 °C for 1 min, continued by 20 cycles of denaturation at 98 °C for 10 s, annealing at 48 °C for 30 s and extension at 72 °C for 3 min and 30 s, followed by a final extension at 72 °C for 7 min. Amplification was checked by electrophoresis in 1% agarose gel. Two PCRs were made per sample (duplicates), mixed and purified using GENECLEAN^®^ Turbo Kit (MP Biomedicals).

### Next-generation sequencing (NGS) of full-length viral genomes

After full-length amplification, a total of 26 cDNA fragments were sequenced: six samples from EU type single infections (three samples from plants grown at 20 °C and other three from plants grown at 30 °C), six samples from CH2 type single infection (three for each temperature), 12 samples from mixed infected plants (three for each isolate and temperature), and also both genomes from the disassembled virions used as starting inoculum were included. Library preparation was performed using the Illumina Nextera XT library preparation kit. Then, libraries were sequenced on the Illumina MiSeq platform, using 250 bp paired-end sequencing reads.

### NGS data analysis

The quality of the raw data was determined using FastQC^56^, and then trimmed by Trimmomatic^57^, which allowed us to remove adapters and trimming low quality bases (Phred score > 30). The quality reads were mapped against the reference genome using bowtie2^58^ and alignments were processed by SAMtools^59^. The SNP calling was performed with different variant callers: LoFreq^60^, FreeBayes^61^ and SNPGenie^62^. Then, we considered as true variants all those with a frequency > 1%, with a minimum raw depth of 3,000 and neglected those variants that were already present in the samples used as starting inocula. Finally, SNPGenie software was also used for estimation of genetic variability indexes such as nucleotide diversity (*π*; mean number of pairwise nucleotide differences per site across the whole genome), and observed heterozygosity (*H*; mean gene diversity at all polymorphic nucleotide sites in the genome).

### Quantitative *in planta* viral infections for fitness estimation

Tomato plants were mechanically inoculated with EU or CH2 types in single and mixed infections (five plants per treatment). This experiment was carried out twice, in a greenhouse under temperature conditions of 20 °C or 30 °C. Seven dpi, total RNA was extracted using Tri-reagent. Viral RNA accumulation was estimated by real-time quantitative PCR (RT-qPCR) using the Power SYBR Green RNA-to-CT 1-Step Kit (Applied Biosystems) and following the manufacturer’s recommendations. RT-qPCR was performed of each sample, considering that in mixed infection treatment two RT-qPCR had to be done per sample, one with EU specific primers and another with those of the CH2^34^. Ten-fold serial dilutions of viral RNA from the disassembled PepMV-Sp13 and -PS5 virions were stocked at −80 °C and used to generate standard curves during the absolute quantification by RT-qPCR.

### Mathematical modelling

Single and mixed infections were investigated by means of mathematical models focusing in the population dynamics of viral strains with temperature-dependent replication rates (see Section S1 in the Supplementary Information (SI)). The models consider exponential replication of the viral strains and also include competition for limited cellular resources (intra-specific competition for single infections and both intra-specific and inter-specific for mixed infections). The model for single infections is given by the logistic equation:

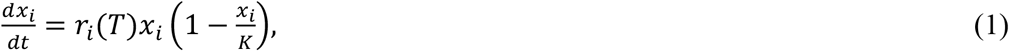

where *x_i_*(t) is the amount of viral RNA of strain *i* ∈ {EU, CH2} at time *t*. For simplicity, EU and CH2 strains will be labeled as E and C, respectively. Here, we assume that the thermal reaction norm for the growth rate *r_i_* (in ng RNA/day) depends on temperature (*T*) as follows:

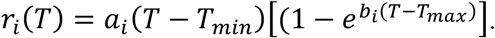

Constants *a_i_* and *b_i_* determine the shape of the reaction norm between a minimum (*T_min_*) and a maximum (*T_max_*) temperatures (for further details on the properties of this reaction norm, see Section S1.1 in the SI). Finally, parameter *K* denotes the maximum amount of viral RNAs that the plants support (carrying capacity in ng of RNA). Equation (1) can be solved analytically to obtain the time-dependent solution:

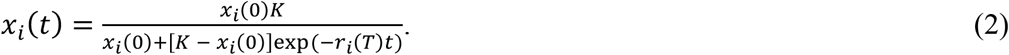

Equation (2) will be used to estimate the parameters for single infections (see below). For the sake of information, the dynamics of Equation (1) is summarized in Section S1.2 in the SI. The model for mixed infections includes the same processes than the one for single infections, incorporating the competition coefficients between the EU and CH2 types, now having a Lotka-Volterra competition model:

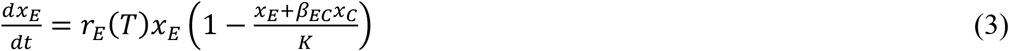

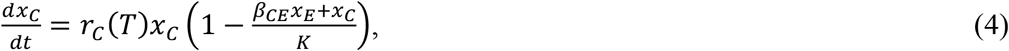

where *β_EC_* measures the intensity of interference that CH2 exerts on EU, and *β_CE_* the complementary (0 < *β_i_* < 1; zero means no interaction and greater values increasing competence). It is not possible to obtain an analytical solution for the two-strains model, and thus the time solutions will be obtained numerically. For the sake of completeness, the dynamics of Equations (3) and (4) are explained in detail in Section 1.3 in the SI.

### Parameters estimation and experimental data fitting

The mathematical models above allowed us to characterize the dynamics of single and mixed infections by considering the main relevant processes of virus replication at the within-host level. Noteworthy, they also allowed to estimate the parameters for each virus strain at different temperatures in both single and mixed infections. Parameter estimations were conducted using an optimization Macroevolutionary Algorithm (MA)^63^. Prior to the analyses with the MA, the experimental data at 7 and 60 dpi were processed to obtain full time series starting at the same initial conditions (Section S2.1 in the SI). The MA is described in detail in Section 2.2 in the SI.

### Statistical analysis

Exploratory statistical analyses were done with SPSS Statistics version 26 (IBM Corp., Armonk, USA) using generalized linear models (GLM). Viral load (*VL*) data were fitted to a model that incorporates three orthogonal factors, one nested factor and one covariable. The orthogonal factors were the type of isolate (*A* ∈ {EU, CH2}), the type of infection (*I* ∈ {single, mixed}) and the experimental temperature (*T* ∈ {20, 30} °C). The five replicate plants (*R* ∈ {1, …, 5}) was nested within the interaction of the three orthogonal factors. Time was used as covariable (*t* ∈ {7, 60} dpi). The full model equation reads:

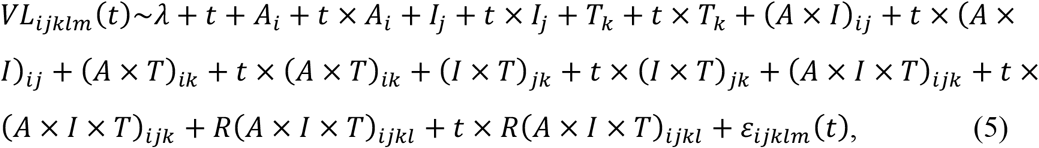

where *λ* corresponds to the grand mean value of *VL* and *ε* represents the sampling error. A Gamma distribution with a log-link function was assumed based on its lowest *BIC* among a set of competing models.

The count of SNPs (*SC*) per sampled population 60 dpi were fitted to a model that incorporates *A*, *I* and *T* as orthogonal factors and uses three replicate plants to estimate the error. In this case, the model equation reads

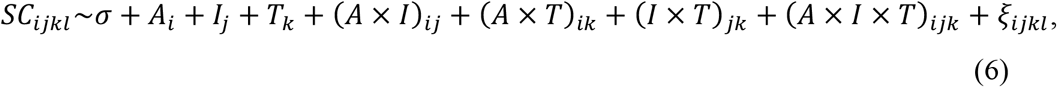

where *σ* corresponds to the grand mean value of *SC* and *ξ* represents the sampling error. A Poisson distribution with log-link function was assumed based on its lowest *BIC* among a set of competing models. The significance of each term in the models was evaluated using a likelihood-ratio test (LRT) which asymptotically follows a *χ^2^* distribution. In addition, the magnitude of the effect associated with each term was evaluated using the 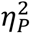 statistic which measures the proportion of total variability in the trait attributable to each factor in the model. Conventionally, values of 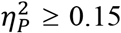 are considered as large effects.

### Data availability

All sequencing information and data that support the findings of this study have been deposited in the NCBI GenBank with the BioProject code PRJNA639566, under accession numbers SRR12017758-SRR12017783. The codes (in C language) developed to study the mathematical models and the MA algorithm are available upon request. Data obtained from all the analyses performed in this work are available as supplementary tables.

## ACKNOWLEDGEMENTS

We thank M.C. Montesinos for technical assistance.C.A. was supported by funding of the Ministry of Economy, Industry and Competitiveness (MINECO, Spain) within a PhD programme grant (FPU16/02569). This work was supported by the Agencia Estatal de Investigaación-FEDER grants AGL2014-59556-R and AGL2017-89550-R to P.G. and PID2019-103998GB-I00 to S.F.E. J.S. has been partially funded by the CERCA Programme of the Generalitat de Catalunya, MINECO grant MTM2015-71509-C2-1-R, Agencia Estatal de Investigación grant RTI2018-098322-B-100, and by a Ramón y Cajal contract (RYC-2017-22243).

Supplementary Information accompanies the paper on the Journal’s website.

## SUPPLEMENTARY INFORMATION

**Table S1.** Nucleotide variants (SNPs), position, frequency and type of change in the PepMV genome for each replicate in single and mixed infection type at 20 °C and 30 °C temperature conditions.

**Table S2.** Data of the experiments (single and mixed infections at 20 °C and 30 °C for 7 and 60 dpi): raw and processed data.

**Table S3.** Statistics obtained with the Macroevolutionary Algorithm for parameters optimization for single infections: estimation of replication rates and carrying capacities for EU and CH2 strains at 20 °C and 30 °C.

**Table S4.** Statistics obtained with the Macroevolutionary Algorithm for parameters optimization for mixed infections: estimation of competition coefficients and carrying capacities.

