## Supplemental material for "Increasing growth temperature alters the within-host competition of viral strains and influences virus genetic variation"

Supplementary Information

Cristina Alcaide[1], Josep Sardanyés[2,3,4], Santiago F. Elena[5,6], Pedro Gómez[1,*]

[1]Centro de Edafología y Biología Aplicada del Segura (CEBAS)- CSIC, Departamento de Biología del Estrés y Patología Vegetal, PO Box 164, 30100, Murcia, Spain

[2]Centre de Recerca Matemàtica (CRM), [3] Barcelona Graduate School of Mathematics and [4]Dynamical Systems and Computational Virology, Associated Unit Instituto de Biología Integrativa de Sistemas (I$^2$SysBio) - CRM. Edifici C, Campus de Bellaterra 08193 Barcelona, Spain

[5]I$^2$SysBio, Consejo Superior de Investigaciones Científicas-Universitat de València, València, Spain

[6]The Santa Fe Institute, Santa Fe, New Mexico, USA.

$*$ corresponding author

**(June 30, 2020)**

### 1 Mathematical models with temperature-dependent virus replication

In the next sections we introduce and analyse the mathematical models used to investigate the dynamics of single and mixed infections at different temperatures. We will refer the viral load (variable $x_i$ in the models) of each strain using as $i$ subindices $E$ for the European (EU) and $C$ for the Chilean (CH2) strains of pepino mosaic virus (PepMV).

#### 1 Modelling the effect of temperature in virus replication (thermal reaction norm)

In this section we introduce the function to model temperature ($T$)-dependent replication rates (thermal reaction norm) of viral gRNA, dubbed $r_i(T)$ with $i \in \{E, C\}$. This function (see Refs. [1, 2, 3] for details) reads:

$$r_i(T) = a_i(T - T_{min})\Big[1 - \exp(b_i(T - T_{max}))\Big]. \tag{S.1}$$

Equation (S.1) produces a "bell-shape" function and assumes that gRNA virus replication increases as temperature grows from a minimum temperature, diminishing when temperature becomes too high. Parameters $a_i$ and $b_i$ are a-dimensional i.e., they have no units, and allow to tune the height and shape of the bell. The units for the replication rates $R_i(T)$ are ng gRNA/day. In our analyses we will set the range of temperatures between $T_{min} = 15\,^oC \leq T \leq 50\,^oC = T_{max}$. To illustrate the behaviour of Eq. (S.1) we show, in Fig. S1, the values of replication within this chosen range of temperatures. Different values of $a_i$ are displayed in the panels, each of them including different values of $b_i$. These two constants, $a_i$ and $b_i$, will be estimated from the experimental data for single infections to determine the replication rates of each strain at the two studied temperatures of 20 $^o$C and 30 $^o$C, thus obtaining the thermal reaction norms for each PepMV strain..

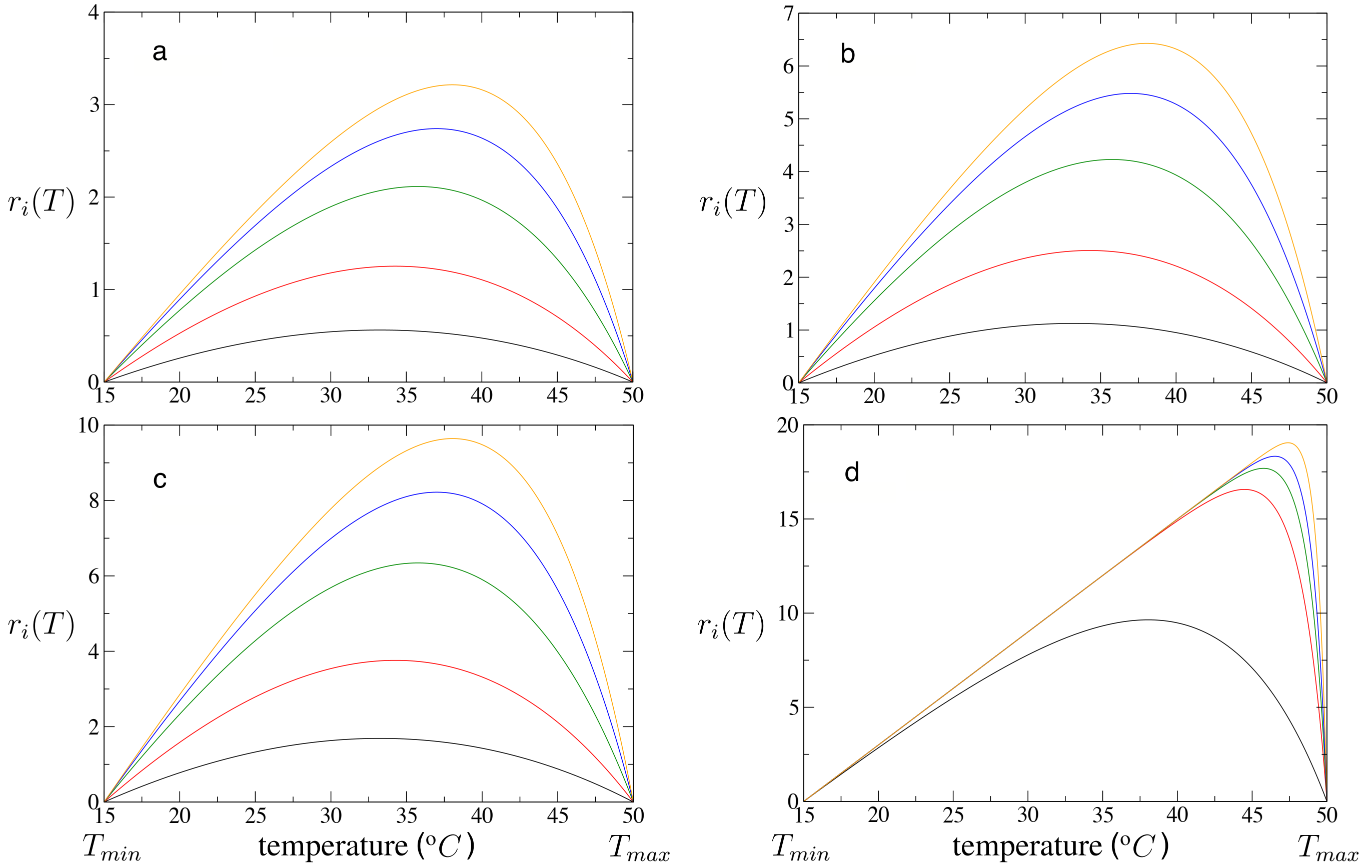

Figure S1: Shape of the reaction norms given by the replication rates, $r_i(T)$ (see Eq. (S.1)) as a function of the temperature, $T$, for different values of parameters $a_i$ and $b_i$. We show results for $a_i = 0.2$ (a), $a_i = 0.4$ (b), $a_i = 0.6$ (c), using different values of $b_i$: 0.010 (black), 0.025 (red), 0.050 (green), 0.075 (blue), and 0.100 (orange). Panel (d) has been computed setting $a_i = 0.6$ and using larger values of $b_i$: 0.10 (black), 0.50 (red), 0.75 (green), 1.00 (blue), and 1.50 (orange).

Finally, let us show that $r_i(T)$ is either zero or positive within the range $T_{min} \leq T \leq T_{max}$, for $a_i$, $b_i > 0$. Let us denote $\alpha = (T - T_{min})$. It is clear that $\alpha = 0$ if $T = T_{min}$ and $\alpha > 0$ if $T > T_{min}$. Also, let us define $\gamma = b_i(T - T_{max})$. Notice that $\gamma = 0$ for $T = T_{max}$, and thus $r_i(T) = 0$. The other case, $T < T_{max}$, involves $\gamma < 0$ and thus the term $\exp(\gamma)$ in Eq. (S.1) is lower than 1, involving $r_i(T) > 0$ for $T_{min} < T < T_{max}$.

#### 2 Mathematical model for single infections: dynamics

The dynamics of virus replication for single infections is described by the next one-variable model::

$$f(x_i) = \frac{dx_i}{dt} = r_i(T)\, x_i \left(1 - \frac{x_i}{K}\right). \tag{S.2}$$

Here, variable $x_i$ is the viral load of each strain $i \in \{E, C\}$, EU: subindex E; CH2: subindex C. This equation is the well-known logistic model, which includes the temperature-dependent growth rate (reaction norm function). The growth rate is exponential for small population sizes and the virus populations will grow until the carrying capacity, $K$ (ng gRNA) is achieved. This can be easily shown by means of linear stability analysis (see below). Actually, this system can be solved analytically, giving as a solution:

$$x_i(t) = \frac{x_i(0)\, K}{x_i(0) + (K - x_i(0)) \exp(-r_i(T)\, t)}, \tag{S.3}$$

where $x(0)$ is the initial condition and $t$ time. Despite having an explicit time-solution and being a very well-known model, let us remind the dynamical properties of the logistic model, now considering the temperature-dependent replication rate. This system has two equilibrium points, namely: $P_1^* = 0$, and $P_2^* = K$. The stability of this equilibrium is determined from

$$\lambda = \frac{df(x_i)}{dx_i} = r_i(T) \left(1 - \frac{2\, x_i}{K}\right).$$

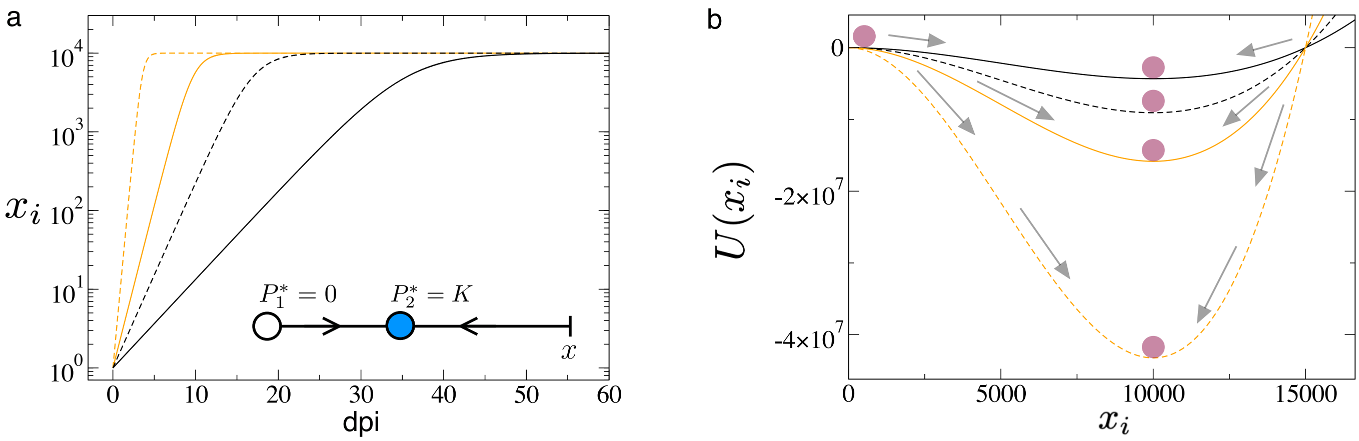

Figure S2: (a) Dynamics obtained from Eq. (S.3) for 60 days post-inoculation (dpi) and using the values of Fig. S2(a), with $a_i = 0.2$, $b_i = 0.01$ (black) and $b_i = 0.1$ (orange), and $K = 10^4$ (shown in linear-log scale). As initial condition we have used $x_i(0) = 1$. Solid and dashed lines correspond to temperatures of 20 ºC and 30 ºC, respectively. The inset displays the phase space of Eq. (S.2), with two equilibrium points, $P_1^*$ (white circle) and $P_2^*$ (blue circle). The arrows indicate the stability of both equilibria. (b) Potential function obtained from Eq. (S.4) using the same parameter values than in panel (a) and represented with same colours and line styles. The local minimum corresponds to the equilibrium point $P_2^* = K$, which is stable.

It is easy to show that $P_1^*$ is unstable since $\lambda(P_1^*) = r_i(T) > 0$, for $T_{min} < T < T_{max}$. Also $\lambda(P_2^*) = -r_i(T) < 0$, and thus $P_2^*$ is an attractor for $T_{min} < T < T_{max}$. That is, for any initial condition, the virus population will always achieve the carrying capacity. Here, the equilibrium values do not depend on $r_i(T)$. We note that the carrying capacity of plant cells may do vary depending on the temperature conditions, thus different equilibria may be achieved at different temperatures (as observed in the experiments). The values of $K$ will be estimated from the experiments (see Section 2 below). Parameter $r_i(T)$ has an important role in the transient towards the equilibrium. That is, different temperatures will make the system to achieve faster or slower the carrying capacity.

Finally, one can easily prove that the system will always achieve the equilibrium point $P_2^*$ for $r_i(T) > 0$. This can be done by taking the limit of Eq. (S.3) as $t \to +\infty$, having:

$$\lim_{t \to +\infty} x_i(t) = \frac{x_i(0)\,K}{x_i(0) + (K - x_i(0))\exp(-r_i(T)\,t)} = K.$$

The time dynamics of Eq.(S.2) can be visualized in Fig. S3(a) by means of time series, obtained from the explicit solution given by Eq. (S.3). The inset in this figure displays a schematic diagram of the phase space, with the equilibria $P_1^*$ being unstable and $P_2^*$ stable. Here, as shown with the stability analysis, the system achieves the carrying capacity at equilibrium. As mentioned, temperature has an important effect at the level of transients, since how fast the equilibrium is approached depends on the replication rate of the viral gRNA. For the parameter values chosen in Fig. S3(a) $x_i$ achieves equilibrium faster at 30 ºC (dashed time series) than at 20 ºC (solid time series). Another way of visualising the dynamics for one-variable systems is by representing a potential function, computed from:

$$U(x_i) = -\int f(x_i)\,dx_i = r_i(T)\,x_i^2\left(\frac{x_i}{3K} - \frac{1}{2}\right). \qquad \text{(S.4)}$$

The potential is a cubic polynomial function and appears parabolic in the studied range of $x_i$ in Fig. S3(b), with a minimum at the equilibrium value $P_2^* = K = 10^4$. Notice that the wells are deeper at 30 ºC, meaning that the equilibrium point is much more attracting. This feature also explains why the equilibrium is achieved faster compared to the dynamics at 20 ºC.

#### 3 Mathematical model for mixed infections: dynamics and transitions

The dynamics for mixed infections is similar to the one used for single infections, considering temperature (T)-dependent replication rates, and including competition between both strains. The strength of competition of strain $j$ on species $i$ is introduced

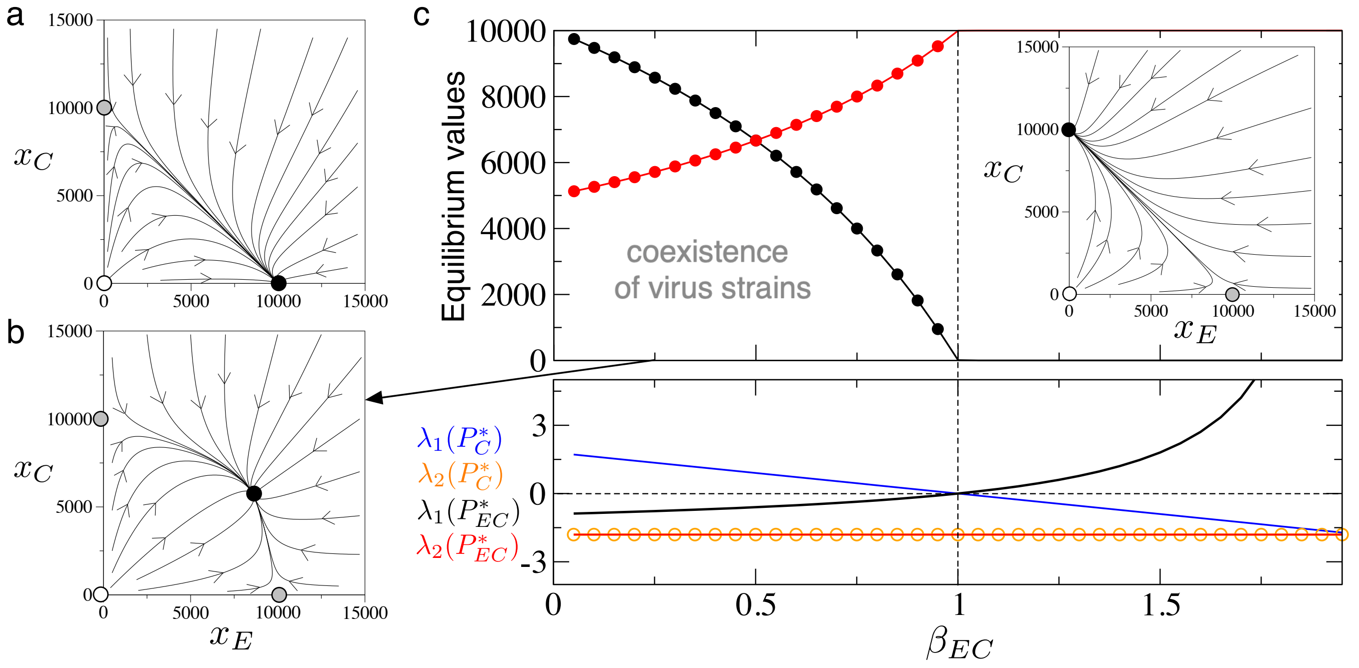

Figure S3: Dynamics of Eqs. (S.5)-(S.6). (a) Phase portrait showing the outcompetition of $x_C$ by $x_E$, with $\beta_{EC} = 0.5$ and $\beta_{CE} = 1.5$ (the arrows indicate the direction of the orbits; equilibrium points are shown with solid circles: repeller (white), saddle point (grey), attractor (black)). (b) Coexistence dynamics, observed in the experiments with mixed infections, here with $\beta_{EC} = 0.25$ (value marked with the arrow in the bifurcation diagram in (c)). (c) Bifurcation diagram increasing the competition strength of $x_C$ on $x_E$, computed with $0.05 \leq \beta_{EC} \leq 1.95$ and $\beta_{CE} = 0.5$. Notice that the two strains coexist for $0 < \beta_{EC} < 1$. The solid line and the solid dots in black indicate the equilibrium values for $x_E$ obtained numerically and analytically, respectively. In red we show the same results for $x_C$. At $\beta_{EC} = 1$ the integRNAl stable point collides with the fixed point $P_C^*$ in a transcritical bifurcation. For $\beta_{EC} > 1$, the equilibrium $P_C^*$ becomes stable. The lower panel displays the eigenvalues for the fixed points $P_C^*$ and $P_{EC}^*$. In all the analyses: $a_E = a_C = 0.8$, $b_E = b_C = 0.02$, $K = 10^4$, and $T = 20\,^{\mathrm{o}}C$.

with (a-dimensional) parameters $\beta_{ij}$, $i, j \in \{E, C\}$, $i \neq j$. We use the Lotka-Volterra competition model, given by:

$$\frac{dx_E}{dt} = r_E(T)\,x_E\left(1 - \frac{x_E + \beta_{EC}\,x_C}{K}\right), \tag{S.5}$$

$$\frac{dx_C}{dt} = r_C(T)\,x_C\left(1 - \frac{\beta_{CE}\,x_E + x_C}{K}\right). \tag{S.6}$$

Equations (S.5)-(S.6) have four different equilibrium points, namely: $P_0^* = (0,0)$, $P_E^* = (K,0)$, $P_C^* = (0,K)$, and $P_{EC}^* = (x_E^*, x_C^*)$, with

$$x_E^* = \frac{K(\beta_{EC} - 1)}{(\beta_{EC}\,\beta_{CE} - 1)}, \qquad x_C^* = \frac{K(\beta_{CE} - 1)}{(\beta_{EC}\,\beta_{CE} - 1)}.$$

Notice that the fixed points are also independent of the replication rates, $r_i(T)$, being determined as well by the carrying capacity and by the competition constants. However, as we will see below, the replication rates also play an important role in the transients towards equilibria.

The stability of the equilibrium points can be studied by means of the eigenvalues of the Jacobian matrix evaluated at the equilibrium. This matrix reads:

$$\mathcal{J} = \begin{pmatrix} r_E(T)(K - 2\,x_E - x_C\,\beta_{EC})/K & -\,r_E(T)\,x_E\,\beta_{EC}/K \\ -r_C(T)\,x_C\,\beta_{CE}/K & r_C(T)(K - 2\,x_C - x_E\,\beta_{CE})/K \end{pmatrix}.$$

The stability of the equilibrium $P_0^*$ is computed from the characteristic equation $\det|\mathcal{J}(P_0^*) - \lambda I| = 0$. The eigenvalues are here given by $\lambda_1(P_0^*) = r_E(T)$ and $\lambda_2(P_0^*) = r_C(T)$. Hence, for positive values of the replication rates this equilibrium point is a repeller.

The eigenvalues of the Jacobian matrix for the equilibrium point $P_E^*$ are $\lambda_1(P_E^*) = -r_E(T)$ and $\lambda_2(P_E^*) = r_C(T)(1-\beta_{CE})$. Notice that the first eigenvalue is negative (when $r_E(T) > 0$) and the stability of this equilibrium depends on the second

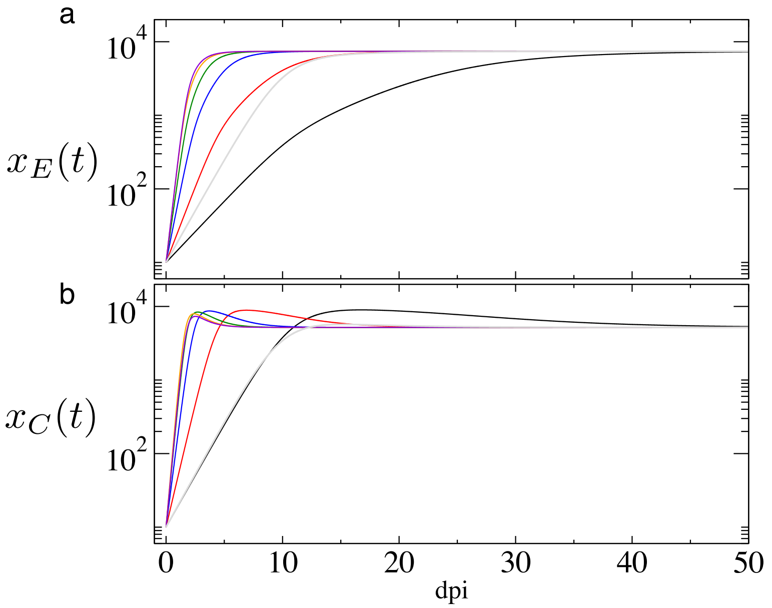

Figure S4: Time dynamics (days post-inoculation (dpi) in linear-log scale) for the viral strains EU ($x_E(t)$ in panel (a)) and CH2 ($x_C(t)$ in panel (b)) at different temperatures: 17$^oC$ (black); 20 $^o$C (red); 25 $^o$C (blue); 30 $^o$C (green); 35 $^o$C (orange); 40 $^o$C (violet); 49 $^o$C (grey). Here we have used: $a_E = 0.2$, $b_E = 0.1$, $\beta_{CE} = 0.65$, $a_C = 0.4$, $b_C = 0.05$, $\beta_{EC} = 0.5$, and $K = 10^4$.

eigenvalue. For $r_{e,c}(T) > 0$ and $\beta_{CE} > 1$ the equilibrium $P_E^*$ is attractor. For $r_{e,c}(T) > 0$ and $\beta_{CE} < 1$ the equilibrium $P_E^*$ is a saddle. The stability of the equilibrium $P_C^*$ is similar to the explained above. Here the eigenvalues are: $\lambda_1(P_C^*) = r_E(T)(1-\beta_{EC})$ and $\lambda_2(P_C^*) = -r_C(T)$. This equilibrium will be stable when $\beta_{EC} > 1$ and $r_E(T) > 0$. It will be a saddle when $\beta_{EC} < 1$ and $r_E(T) > 0$.

The stability of the coexistence point, $P_{EC}^*$, is also obtained from the eigenvalues of $\det|\mathcal{J}(P_{EC}^*) - \lambda I| = 0$. For this case we have:

$$\lambda_{\pm}(P_{EC}^*) = \frac{r_E(T)(1-\beta_{EC}) + r_C(T)(1-\beta_{CE}) \mp \sqrt{(r_E(T)\,\Theta + r_C(T)\Gamma)^2 + 4\,r_E(T) r_C(T)\,\Theta\,\Gamma(\beta_{EC}\beta_{CE}-1)}}{2\,(\beta_{EC}\,\beta_{CE}-1)}.$$

with $\Theta = \beta_{EC} - 1$ and $\Gamma = \beta_{CE} - 1$.

A global picture of the dynamics of the mixed infections model is provided in Fig. S3. Panel (a) displays a scenario of virus outcompetition, where the EU strain outcompetes the CH2 strain. In panel (b) we display the parametric scenario in which both gRNA strains coexist, as observed in the experiments performed in this work. Panel (c) provides a bifurcation diagram, built plotting the equilibrium values of both strains at increasing values of $\beta_{EC}$. Notice that for $\beta_{EC} < 1$ the two viral strains coexist. At $\beta_{EC} > 1$ the coexistence turns into outcompetition of the strain EU by strain CH2 (see the phase portrait in the inset). The panel below the bifurcation diagram shows the eigenvalues for the fixed points $P_C^*$ and $P_{EC}^*$. Both fixed points collide and interchange stability by means of one eigenvalue, in what is known as a transcritical bifurcation. Finally, Fig. S4 shows time series obtained numerically[1] for different temperature values. Here, as shown for the single infection dynamics, temperatures involving higher replication rates of viral gRNA will allow to achieve the equilibrium points earlier. However, for temperatures close to the maximum values (e.g., 40$^oC$ and 49$^oC$) the dynamics slows down again.

### 2 Fitting of the mathematical models to the experimental data: Parameters estimation

#### 1 Experimental data processing

The experimental data will be processed for the fitting and parameter estimation process. We will build three-point time series from the experimental data using the replicates obtained at 7 and 60 dpi for each viral strain, also including the known initial conditions. That is, for each temperature we will build different three-points time series for both single and mixed infections, having at the end 8 different data sets (4 time series for the single infections plus 4 time series for the mixed infections). Since

[1]Numerical simulations have been performed with a fourth-order Runge-Kutta method with a constant time step $\Delta t = 0.05$.

several replicates have been obtained we will apply the optimisation algorithm described in the next section to all replicates simultaneously. In order to merge all this information into a three-point time series we need to re-scale the experimental data since the initial conditions used experimentally at 7 and 60 dpi were different. The data re-scaling will be applied to all data points (including the initial conditions), using:

$$x_i^s(t) = \left(x_i(t) - x_i^{(t)}(0)\right) + 1, \qquad i \in \{E, C\}, \qquad t \in \{0, 7, 60\},$$

where $x_i^s(t)$ is the scaled data at time $t$ and $x_i^{(t)}(0)$ is the initial condition used for the experiments performed performed during times $t$ ($x_i^{(t=7)}(0) = 150$ ng gRNA or $x_i^{(t=60)}(0) = 899$ ng gRNA). By applying this re-scaling we now take into account the net increase of the population starting at the same initial condition, which is set to 1, since $x_i(0) - x_i^{(t)}(0) = 0$.

#### 2 Macroevolutionary algorithm for paramaters estimation

In order to find the values of the model parameters providing the best fitting to the experimental data we will use a macroevolutionary algorithm (MA) [4]. This algorithm considers a population of vectors of parameters to which a fitness value can be attributed depending on how good are these sets of parameters in fitting the mathematical model to the experimental data. This fitness value is usually given by a distance measure between the simulated (i.e., data obtained from the mathematical model) and the data measured in the experiments (here we will use the least-squares, see below). The MA involves a selection of the fittest parameters due to processes of evolution and macro-extinctions. The MA is implemented as follows:

- Define a population, $\Omega(N)$, with $N$ vectors of parameters, $\vec{\nu}_i^{(k)} = (a_i, b_i, \beta_{ij}, \beta_{ji}, K)^{(k)}$, with $i, j \in \{E, C\}$, $i \neq j$; $1 \leq k \leq N$. Note that for single infections $\beta_{ij} = \beta_{ji} = 0$.
- Define exploratory ranges for each parameter. We give the following ranges for the model parameters:

$$a_i, b_i \in [10^{-5}, 10^1], \qquad \beta_{ij}, \beta_{ji} \in (0, 1), \qquad K \in [10^3, 10^{11}]. \tag{S.7}$$

  Since the ranges for parameters $a_i$ and $b_i$ include several orders of magnitude we will sample the exponents from the uniform distribution $U(-5, 1)$. The analyses developed for the mixed infections model (Section 1.3 above) indicated that the values of $\beta_{EC}$ and $\beta_{CE}$ must be positive and lower than 1 to ensure coexistence of strains, as observed in the competition experiments. Since we are interested in the relative values of $\beta_{EC}$ and $\beta_{CE}$, and, due to the dynamical restrictions discussed above, we will explore the range $(0, 1)$ for both competition coefficients. Finally, the carrying capacity will be explored for the range $10^3 \leq K \leq 10^{11}$, also sampling the exponents from the uniform distribution $U(3, 11)$.
- Initialise $\Omega(N, \tau = 0)$ with random values within the intervals defined in (S.7). Once the initial population of $N$ vectors is initialised, the macroevolutionary generations, $\tau$, start. At each generation $\tau$ we apply the next rules:
  1. For each vector $\vec{\nu}_i^{(k)}$ we compute the distance (labeled as $d_i^{(k)}$) between the simulated data, $x_i^{(k)}(t)$, obtained with the vector of parameters $\vec{\nu}_i^{(k)}$, and the re-scaled experimental data, now labeled as $\varepsilon(t)$, by means of the least squares. The simulated solutions for the single infections are computed from the analytical solution given by Eq. (S.3), while for mixed infections the solutions are obtained integrating equations (S.5)-(S.6) numerically[1]. Recall that for both single and mixed infections the initial conditions are $x_i(0) = 1$. Since several replicates are available for each time point ($t_1 = 7$ dpi and $t_2 = 60$ dpi), we will compute a single distance value by means of the least squares, employing:

$$d_i^{(k)} = F \cdot \frac{1}{(n_7 + n_{60})} \left( \sum_{j=1}^{n_7} \left(x_j^{(k)}(t_1) - \varepsilon(t_1)\right)^2 + \sum_{m=1}^{n_{60}} \left(x_m^{(k)}(t_2) - \varepsilon(t_2)\right)^2 \right), \tag{S.8}$$

     $n_7$ and $n_{60}$ being the number of experimental replicas at 7 and 60 dpi, respectively. Due to the large viral loads measured in the experiments we will multiply the distances by a factor $F = 10^{-4}$ to handle lower numbers. This has no effect on the parameters optimisation process since it will be applied to all the simulations. For single infections we will compute $d_i^{(k)}$ for each strain and for each studied temperature (20 ºC and 30 ºC). For mixed infections we need to compute the distance taking into account the amounts of gRNA for the two virus strains simultaneously. For this case, the distance is computed using:

$$D_m^{(k)} = \frac{1}{2}\left(d_E^{(k)} + d_C^{(k)}\right). \tag{S.9}$$

2. Order the whole vectors of the population from lowest to largest distances.
3. Select a 25% of the vectors with the lowest distances, removing all others (macroextinctions).
4. Then, the population $\Omega(N, \tau+1)$ will be initialised with the 25% of the survival vectors (following the order from lower to higher distances). Together with the selected vectors, the population at $\tau+1$ will also include the same selected vectors slightly perturbed. Perturbations consider small variations i.e., $\vec{\nu}_i^{(k)} = (a_i(1 \pm \mu), b_i(1 \pm \mu), K(1 \pm \mu), \beta_{ij}(1 \pm \mu), \beta_{ji}(1 \pm \mu))^{(k)}$. Here we will generally use values of $\mu$ selected uniformly at each time generation within ranges $\mu \in [10^{-12}, 10^{-2}]$. After doing so, the population $\Omega(N, \tau+1)$ will be again composed of $\xi = N/2$ vectors. Then, the remaining and extincted population of vectors $N - \xi$ will be replaced at $\tau+1$ by other vectors taking random values within the ranges given in (S.7).
5. Go to 1.

The previous algorithm involves a selection of the best parameters, including massive extinctions and evolution of the best selected vectors along the iterations of the MA. That is, an evolutionary optimisation of parameters is done. In our simulations we will use a population size of $N = 4,000$ vectors of parameters. At the end of each MA simulation we will obtain a population of $N/4 = 10^3$ optimised vectors of parameters, that will be averaged and considered as best mean values of the estimated parameters. We note that we will also record the parameter vector providing the best fit along the whole MA. Also, we will compute the mean of the parameters providing the best fit for each replica of the MA algorithm. For each experimental data set we will run 50 replicates of the MA. Each replicate involves a total exploration of $4,000$ vectors of parameters (initial population) plus $3,000$ vectors that will be modified (including those with small random perturbations and those extinct) by iteration of the MA algorithm. For single infections we will run the MA over $\tau = 10^7$ generations. This gives a total amount of more than $1.5 \times 10^{12}$ different combinations of explored parameters. For mixed infections, which have a higher computational cost due to the numerical computation of the solutions, we will use set $\tau = 10^6$ generations. For this case we will explore more than $1.5 \times 10^{11}$ different parameter combinations.

### 3 Parameters estimation for single infections

#### 3.1 Data for EU strain replication at 20 ºC

The parameters providing the best fit, with distance $d_E = 168,999,048.553$, are:

$$\begin{aligned} \text{best } a_E &= 1.5519, \\ \text{best } b_E &= 0.8138, \\ \text{best } R_E(T) &= 7.7595 \text{ ng gRNA/day}, \\ \text{best } K &= 1,968,217.8367 \text{ ng gRNA}. \end{aligned}$$

The fitting of the mathematical model to the experimental data using these parameters is displayed in Fig. S5(a) in black.

The mean values ($\pm SD$) hereafter computed from the 50 replicas of the MA of the optimised parameters (obtained at the end of the MA) are:

$$\begin{aligned} \bar{a}_E \pm SD &= 4.2449 \pm 0.1052, \\ \bar{b}_E \pm SD &= 1.5394 \pm 0.1209, \\ \bar{R}_E(T) &= 21.2247 \text{ ng gRNA/day}, \quad (\bar{R}_E(T) \text{ computed from } \bar{a}_E \text{ and } \bar{b}_E \text{ above}), \\ \bar{K} \pm SD &= 1,968,217.8737 \pm 0.2252 \text{ ng gRNA}. \end{aligned}$$

Finally, the averages ($\pm SD$) of the best parameter vectors (hereafter also computed from the 50 replicates) are:

$$\begin{aligned} \langle \text{best } a_E \rangle \pm SD &= 4.0668 \pm 2.2586, \\ \langle \text{best } b_E \rangle \pm SD &= 1.6117 \pm 2.6222, \\ \langle \text{best } R_E(T) \rangle &= 20.3342 \text{ ng gRNA/day}, \quad (\text{computed from } \langle \text{best } a_E \rangle \text{ and } \langle \text{best } b_E \rangle \text{ above}), \\ \langle \text{best } K \rangle \pm SD &= 1,968,217.8559 \pm 0.0232 \text{ ng gRNA}. \end{aligned}$$

The fittings of the mathematical model to the experimental data for the mean values of both optimised parameters and best vectors of parameters are shown in Fig. S6(a) in black for the experimental replicates (upper panel) and their mean values (lower panel).

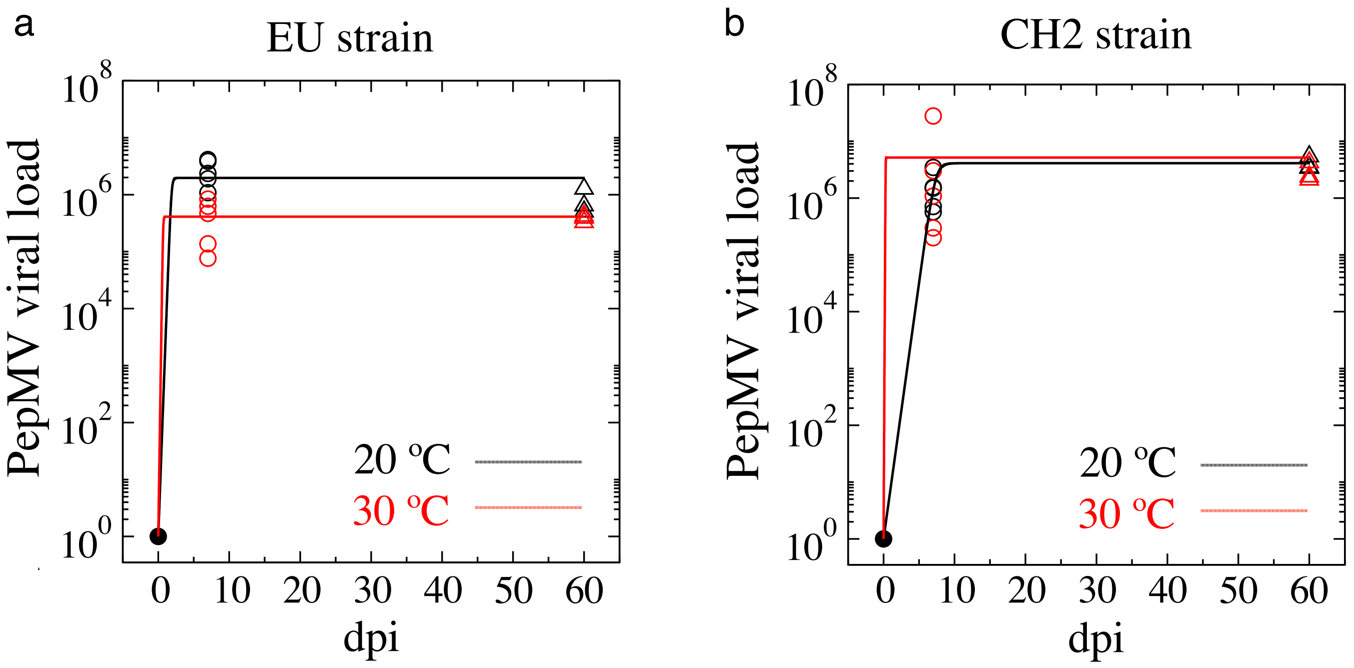

Figure S5: Fitting of the mathematical model to the experimental data using the best vector of parameters obtained from 50 replicates of the Macroevolutionary Algorithm (MA) for single infections for EU (a) and CH2 (b) strains. Open circles show experimental data at 20 ºC (black) and 30 ºC (red); time trajectories show the dynamics obtained with the mathematical model; solid dots show the initial conditions. The fittings have been performed using the explicit solution given by Eq. (S.3).

##### 3.2 Data for EU strain replication at 30 ºC

The parameters providing the best fit, now with distance $d_E = 5,316,845.058$, are:

$$\begin{aligned} \text{best } a_E &= 1.2681, \\ \text{best } b_E &= 0.3841, \\ \text{best } R_E(T) &= 19.0123 \text{ ng gRNA/day}, \\ \text{best } K &= 411,543.4159 \text{ ng gRNA}. \end{aligned}$$

The fitting of the model to the data obtained with these parameters is displayed in Fig. S5(b) in red.

The averages ($\pm SD$) of the optimised parameters here are:

$$\begin{aligned} \bar{a}_E \pm SD &= 3.1413 \pm 0.1074, \\ \bar{b}_E \pm SD &= 1.4630 \pm 0.1007, \\ \bar{R}_E(T) &= 47.1194 \text{ ng gRNA/day}, \quad (\bar{R}_E(T) \text{ computed from } \bar{a}_E \text{ and } \bar{b}_E \text{ above}), \\ \bar{K} \pm SD &= 411,543.4072 \pm 0.0245 \text{ ng gRNA}. \end{aligned}$$

The averages ($\pm SD$) of the best parameter vectors are:

$$\begin{aligned} \langle \text{best } a_E \rangle \pm SD &= 3.7723 \pm 3.0843, \\ \langle \text{best } b_E \rangle \pm SD &= 2.5047 \pm 3.0466, \\ \langle \text{best } R_E(T) \rangle &= 56.5843 \text{ ng gRNA/day}, \quad (\text{computed from } \langle \text{best } a_E \rangle \text{ and } \langle \text{best } b_E \rangle \text{ above}), \\ \langle \text{best } K \rangle \pm SD &= 411,543.4346 \pm 0.1416 \text{ ng gRNA}. \end{aligned}$$

The fittings of the mathematical model to the experimental data for the mean values of both optimised parameters and best vectors of parameters are shown in Fig. S6(a) in red for the experimental replicates (upper panel) and their mean values (lower panel).

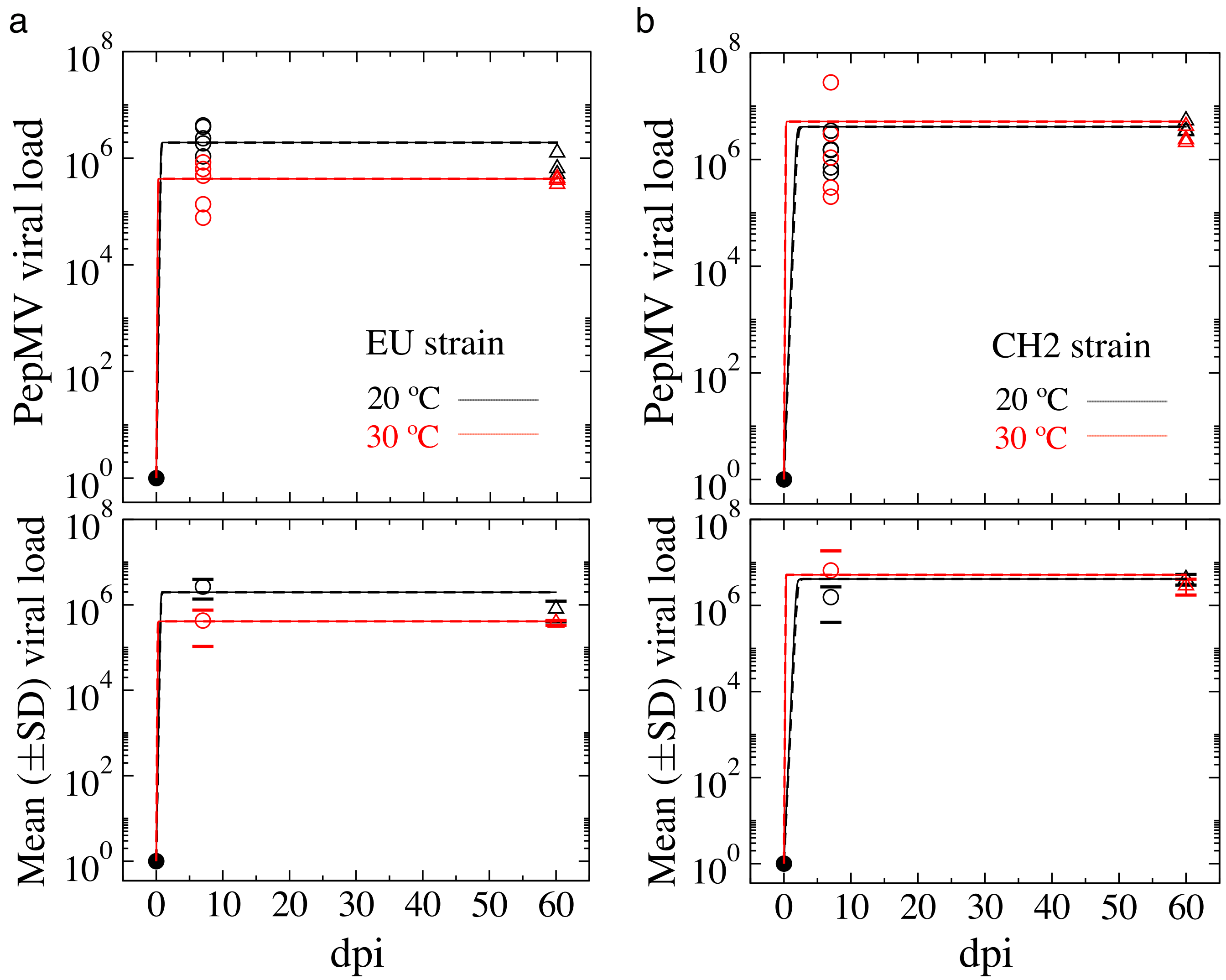

Figure S6: Fitting of the mathematical model to the experimental data for single infections for EU (a) and CH2 (b) strains. Open circles show experimental data at 20 ºC (black) and 30 ºC (red); time trajectories show the dynamics obtained with the mathematical model; solid dots show the initial conditions. Upper and lower plots display total and mean (± SD) viral load over time, respectively. The fittings have been performed using the explicit solution given by Eq. (S.3). All plots display the fitting with the mean values of the optimised parameters (from 50 replicates, solid line); and the mean values of the best vectors (from the 50 replicates, dashed lines). Notice that the time series appear overlapped.

##### 3.3 Data for CH2 strain replication at 20 ºC

The vector of parameters providing the best fit (distance $d_C = 98,402,070.291$), are:

$$\begin{aligned} \text{best } a_C &= 1.1382, \\ \text{best } b_C &= 0.0151, \\ \text{best } R_C(T) &= 2.0757 \text{ ng gRNA/day}, \\ \text{best } K &= 4,110,229.0092 \text{ ng gRNA}. \end{aligned}$$

The fitting of the mathematical model to the experimental data using these parameters is displayed in Fig. S5(b) in black.

Mean values ($\pm SD$) of the optimised parameters are given by:

$$\begin{aligned}\bar{a}_C \pm SD &= 1.6448 \pm 1.4612,\\ \bar{b}_C \pm SD &= 0.8539 \pm 1.0798,\\ \bar{R}_C(T) &= 8.2238 \text{ ng gRNA/day}, \quad (\bar{R}_C(T) \text{ computed from } \bar{a}_C \text{ and } \bar{b}_C \text{ above}),\\ \bar{K} \pm SD &= 4,110,226.7122 \pm 10.4508 \text{ ng gRNA}.\end{aligned}$$

Averages ($\pm SD$) of the best parameter vectors:

$$\begin{aligned}\langle\text{best } a_C\rangle \pm SD &= 1.4879 \pm 1.6924,\\ \langle\text{best } b_C\rangle \pm SD &= 0.9969 \pm 1.9187,\\ \langle\text{best } R_C(T)\rangle &= 7.4398 \text{ ng gRNA/day}, \quad (\text{computed from } \langle\text{best } a_C\rangle \text{ and } \langle\text{best } b_C\rangle \text{ above}),\\ \langle\text{best } K\rangle \pm SD &= 4,110,232.8545 \pm 43.4579 \text{ ng gRNA}.\end{aligned}$$

The fittings of the mathematical model to the experimental data for the mean values of both optimised parameters and best vectors of parameters are shown in Fig. S6(b) in black for the experimental replicates (upper panel) and their mean values (lower panel).

##### 3.4 Data for CH2 strain replication at 30 °C

The parameters giving the best fit ($d_C = 7,553,909,950.168$) are given by:

$$\begin{aligned}\text{best } a_C &= 3.5948,\\ \text{best } b_C &= 1.3388,\\ \text{best } R_C(T) &= 53.9226 \text{ ng gRNA/day},\\ \text{best } K &= 5,159,810.7345 \text{ ng gRNA}.\end{aligned}$$

The fitting of the mathematical model to the experimental data using these parameters is displayed in Fig. S5(b) in red. Mean values ($\pm SD$) of the optimised parameters:

$$\begin{aligned}\bar{a}_C \pm SD &= 3.2705 \pm 0.1284,\\ \bar{b}_C \pm SD &= 1.4583 \pm 0.1155,\\ \bar{R}_C(T) &= 49.0568 \text{ ng gRNA/day}, \quad (\bar{R}_C(T) \text{ computed from } \bar{a}_C \text{ and } \bar{b}_C \text{ above}),\\ \bar{K} \pm SD &= 5,159,810.5657 \pm 0.2940 \text{ ng gRNA}.\end{aligned}$$

Averages ($\pm SD$) of the best parameter vectors:

$$\begin{aligned}\langle\text{best } a_C\rangle \pm SD &= 3.6582 \pm 2.8391,\\ \langle\text{best } b_C\rangle \pm SD &= 1.2363 \pm 2.1985,\\ \langle\text{best } R_C(T)\rangle &= 54.8726 \text{ ng gRNA/day}, \quad (\langle\text{best } R_C(T)\rangle \text{ computed from } \langle\text{best } a_C\rangle \text{ and } \langle\text{best } b_C\rangle \text{ above}),\\ \langle\text{best } K\rangle \pm SD &= 5,159,810.7453 \pm 0.0503 \text{ ng gRNA}.\end{aligned}$$

The fittings of the mathematical model to the experimental data for the mean values of both optimised parameters and best vectors of parameters listed above are shown in red in Fig. S6(b) for the experimental replicates (upper panel) and their mean values (lower panel).

Figure S7 shows how the optimisation of the parameters takes place along the MA. The mean distances $\langle d_i \rangle^-$ (plotted extracting an integer number, see below) between the experimental and simulated data show a decrease during the MA generations (in linear-log axes) for: EU strain at 20 °C (a) and 30 °C (b); CH2 strain at 20 °C (c) and 30 °C (d). These distances correspond to the mean values of the first $N/4$ parameters vectors, which will have lower values since they are the selected ones generation after generation. In all of the analyses the mean distances initially show a sharp decline and then a slower decrease towards lower stationary values. The insets display the distances each time a parameter set providing the best fit is found. These improvements are displayed as consecutive events (improvement event) for the sake of clarity. It does not mean that they appear consecutively along the MA. For the sake of visualisation, the values of the distances are plotted extracting, for each replicate of the same experiment, the same integer value to get smaller numbers. This extraction has been only performed to show the decrease of the distances due to the selection of the parameters during the MA in Fig. S7 (and Fig. S10 for mixed infections) with smaller numbers. The sharp declines in the inset of Fig. S7(c) are due to the presence of negative distances for some replicates due to this extraction. We want to emphasise that the comparison of the distances (errors) between experiments for single infections (mixed infections) must be done with the values of $d_i$ ($D_m$).

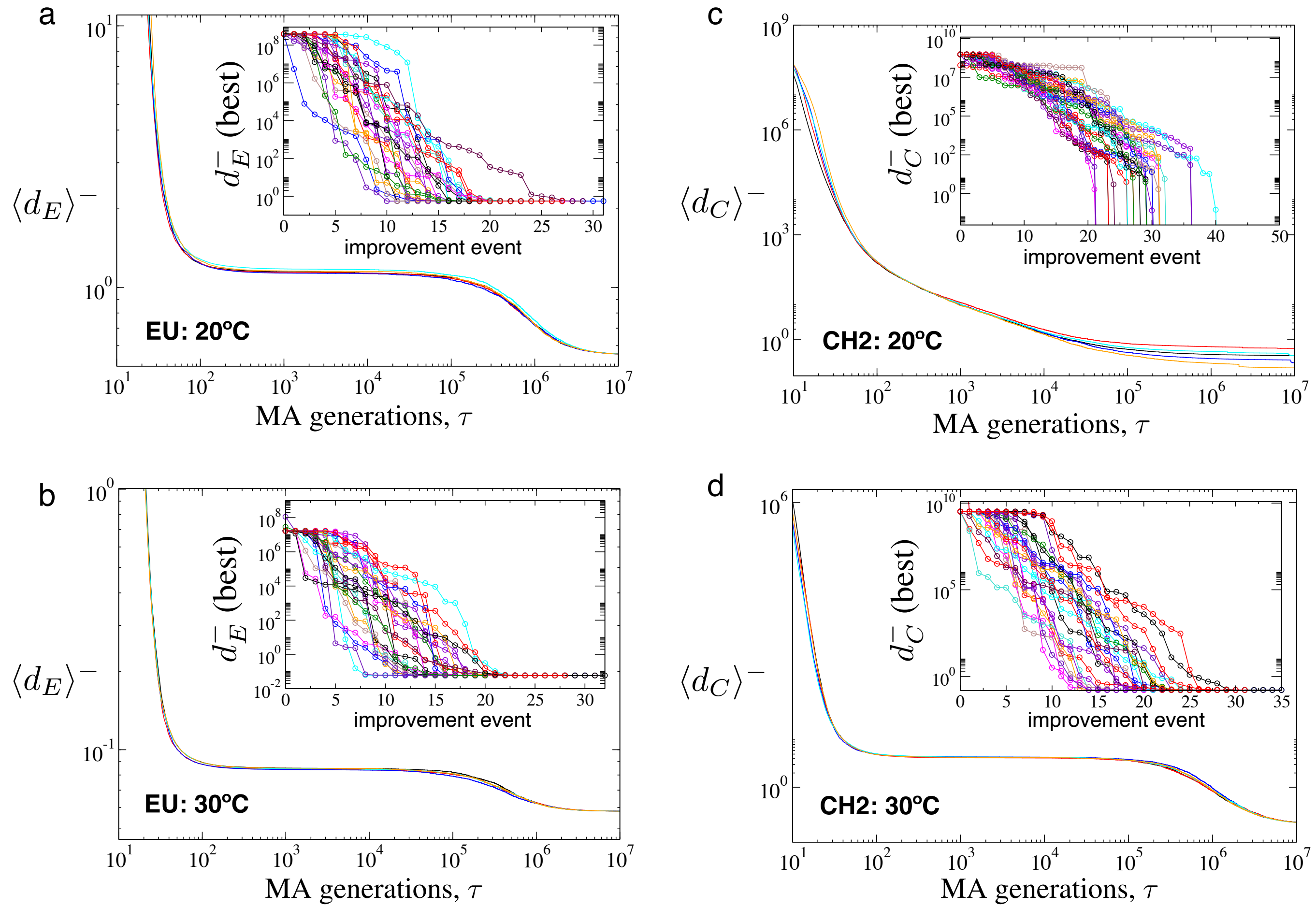

Figure S7: Evolution of the mean distances (shown as $\langle d_i \rangle^-$, see the text), computed from the best $N/4$ vectors of parameters during the optimisation carried out by the Macroevolutionary Algorithm for single infections with strains EU (panel (a) at 20 ºC; panel (b) at 30 ºC) and CH2 (panel (c) at 20 ºC; panel (d) at 30 ºC). Here we show overlapped the results for 5 replicates. The insets display the decrease of the distances for the best vectors of parameters at each improvement event, shown overlapped for 25 replicates.

##### 3.5 Thermal reaction norms for EU and CH2 strains

The estimation of parameters $a_i$ and $b_i$ from the single infections experiments allows the reconstruction of the reaction norm for each strain as a function of temperature $T$ for each virus strain. According to the experimental data and to the chosen function to define the $T$-dependent replication rate, $r_i(T)$, given by Eq. (S.1), replication rates for both strains follow a similar replication profile as a function of $T$. The increase of replication rates with $T$ follows a linear behaviour until $T$ gets close to the maximum value, experiencing a sharp decline. These values shall be taken with caution since they are function-dependent and the replication of the viruses at extreme temperatures (especially at high temperatures) may vary due to physiological changes in plant cells as well as to different enzymatic activities of molecular components crucial for gRNA amplification and virus assembly. For intermediate values, probably, the replication values may follow this linear fashion, but this is pure speculative. Figure S8 displays these results for strains EU (a) and CH2 (b). Each plot shows 6 predictions: 3 for the experiments at 20 ºC (solid lines) and 3 for the ones performed at 30 ºC (dashed lines). These 3 curves represent the shape of the curves computed with the mean values of $a_i$ and $b_i$ (black) obtained at the end of the MA algorithm and averaged over the 50 replicates. The green curves display the profiles obtained from the $a_i$ and $b_i$ values for the best vector of parameters obtained from the full 50 replicas of the MA. Finally, the blue lines show the profiles for the values of $a_i$ and $b_i$ averaged over the best vectors obtained for each replicates of the MA. Despite the curves appear different for some chosen values of $a_i$ and $b_i$ (especially for the best vector of CH2 at 30 ºC), all of them have the same shape, meaning that this result, despite the limitations of the experimental data and the selected function, $r_i(T)$, remains consistent. Future research may experimentally check whether virus replication falls into this type of curves at increasing temperature.

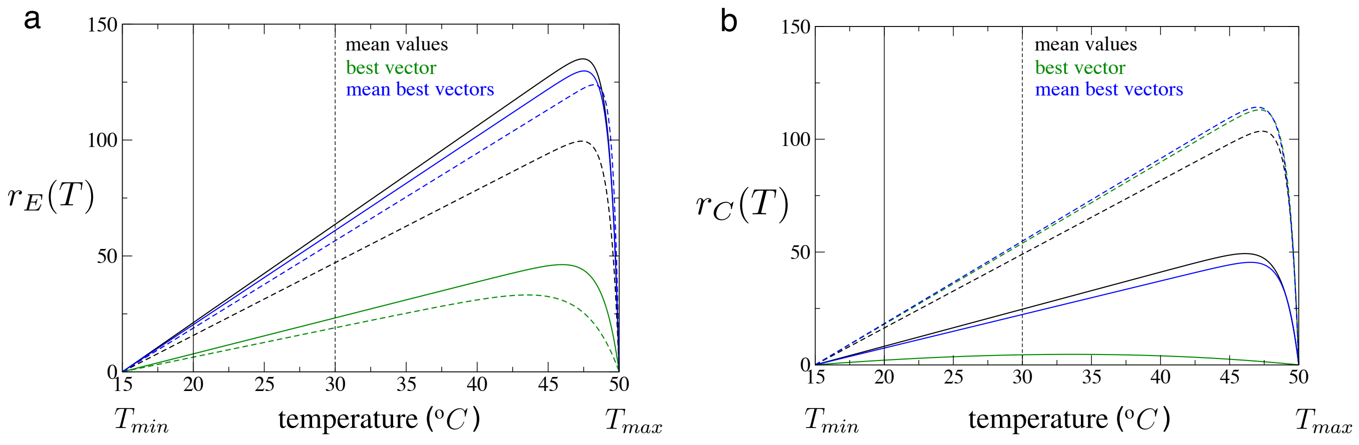

Figure S8: Reaction norms for each of the virus strains at increasing temperatures obtained from the parameters estimated from the experimental data of Section 3. Curves $r_E(T)$ for EU (a) and $r_C(T)$ for CH2 are built from the mean values of the estimated parameters (black) from the 50 replicas of the MA, the best vectors (green); and the mean of the best vectors (blue) also computed from the replicas of the MA. Solid and dashed lines show the data for the experiments at 20 °C and 30 °C, respectively.

#### 4 Parameters estimation for mixed infections

In the previous section we have estimated the parameter values for the single infections independently. This has served us to obtain the parameters concerning the temperature-dependent replication rates and the carrying capacities for each viral strain at each temperature of study. The main goal of the experiments with the mixed infections is to characterise the competition dynamics between EU and CH2 strains at different temperatures, $\beta_{EC}$ and $\beta_{CE}$ being the competition coefficients. To do so we will fix the values of $R_E(T)$ and $R_C(T)$ using the constants $a_E$, $b_E$, and $a_C$, $b_C$ that gave the best fits in the single infections experiments. Since now the two viral strains are co-infecting the same cells, the values of the carrying capacities estimated in the single infections might differ from the ones of mixed ones. Hence, we will introduce as free parameters the vector for the mixed infections $\vec{\nu}_m^{(k)} = (\beta_{EC}, \beta_{CE}, K)^{(k)}$. We emphasise that the competition terms as are introduced in the mathematical model affect the net replication rate of the viral strains and, as mentioned, in order to obtain clear results about the interference due to competition, we here use $R_E(T = 20) = 7.7594$, $R_C(T = 20) = 2.0757$, $R_E(T = 30) = 19.0122$, and $R_C(T = 30) = 53.9225$.

##### 4.1 Data for competition experiments at 20 °C

Fitting with the best fit vector ($d_m = 1,311,471,777.870$), given by:

$$\begin{aligned} \text{best } \beta_{EC} &= 0.9319, \\ \text{best } \beta_{CE} &= 0.0793, \\ \text{best } K &= 7,041,809.0175 \text{ ng gRNA.} \end{aligned}$$

The fitting of the mathematical model with competition to the experimental data using these parameters is displayed in Fig. S9(a), with dynamics for EU and CH2 strains represented in black and green, respectively..

Mean parameter values ($\pm SD$) after the optimisation process:

$$\begin{aligned} \bar{\beta}_{EC} \pm SD &= 0.9319 \pm 1.8 \times 10^{-7}, \\ \bar{\beta}_{CE} \pm SD &= 0.0793 \pm 5.9 \times 10^{-8}, \\ \bar{K} \pm SD &= 7,041,819.8047 \pm 13.2831 \text{ ng gRNA.} \end{aligned}$$

Averages ($\pm SD$) of the better vectors of parameters:

$$\begin{aligned} \langle \text{best } \beta_{EC} \rangle \pm SD &= 0,9318 \pm 1.4 \times 10^{-6}, \\ \langle \text{best } \beta_{CE} \rangle \pm SD &= 0.0793 \pm 4.9 \times 10^{-7}, \\ \langle K \rangle \pm SD &= 7,041,812.2274 \pm 60.9229 \text{ ng gRNA.} \end{aligned}$$

The fittings of the mathematical model to the experimental data for the mean values of both optimised parameters and best vectors of parameters listed above are shown in red in Fig. S10(a) for the experimental replicates (upper panel) and their mean values (lower panel).

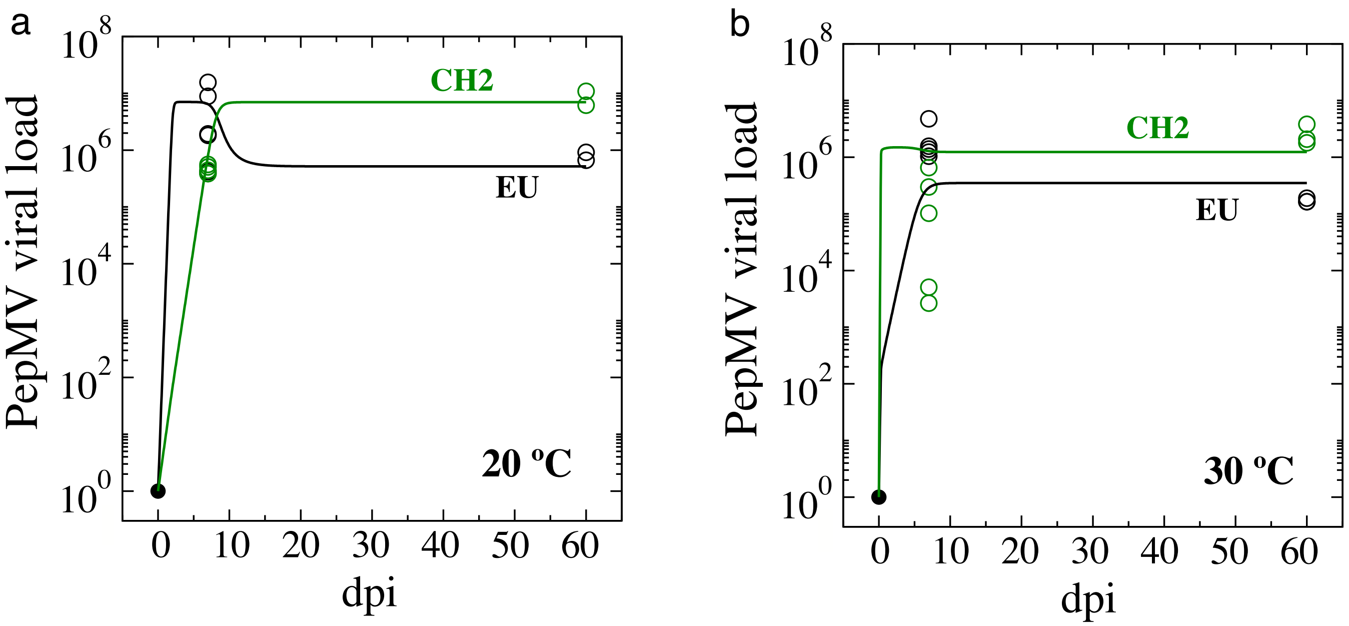

Figure S9: Fitting of the experimental data (open circles) for mixed infections (EU strain (black), CH2 strain (green)) obtained with the mathematical model given by Eqs. (S.5)-(S.6) using the parameters obtained with the best fit from 50 replicas of the MA. Panels (a) and (b) display the dynamics of competition at $T = 20\,^{o}C$ and $T = 30^{o}C$, respectively.

##### 4.2 Data for competition experiments at 30 ºC

Here, due to the large replication rate of CH2 at 30 ºC (i.e., $R_C(T = 30) = 53.9225$) and due to the dispersion of the data (especially of strain CH2 at 7 dpi) we restricted the exploration of $\beta_{EC}$ and $\beta_{CE}$ to obtain better fittings, focusing on good matches between the experimental data and the values obtained with the mathematical model at 60 dpis. The criterium to choose the ranges of the inter-specific competition coefficients has been established by generating multiple time series using the fixed replication rates and tuning $\beta_{EC}$, $\beta_{CE}$, in several pre-defined and narrower ranges. By doing so we observed that better fittings were obtained within the ranges $\beta_{EC} \in [0.9, 1)$ and $\beta_{CE} \in (0, 0.75]$ (results not shown). The resulting mean values ($\pm SD$) of the optimised parameters within these ranges are: The parameters giving the best fit ($d_m = 236,828,771.245$) are:

$$\begin{aligned} \text{best } \beta_{EC} &= 0.9291, \\ \text{best } \beta_{CE} &= 0.7497, \\ \text{best } K &= 1,496,690.8362 \text{ ng gRNA}. \end{aligned}$$

The fitting of the mathematical model with competition to the experimental data using the parameters above is shown in Fig. S9(b), here also with dynamics for EU and CH2 strains represented in black and green, respectively..

$$\begin{aligned} \bar{\beta}_{EC} \pm SD &= 0.9290 \pm 4.16 \times 10^{-5}, \\ \bar{\beta}_{CE} \pm SD &= 0.7490 \pm 7.37 \times 10^{-4}, \\ \bar{K} \pm SD &= 1,494,856.9567 \pm 982.4729 \text{ ng gRNA}. \end{aligned}$$

Finally, the averages ($\pm SD$) of the best vectors obtained during the optimisation are::

$$\begin{aligned} \langle \text{best } \beta_{EC} \rangle \pm SD &= 0.9291 \pm 4.17 \times 10^{-5}, \\ \langle \text{best } \beta_{CE} \rangle \pm SD &= 0.7490 \pm 7.38 \times 10^{-4}, \\ \langle K \ \rangle \pm SD &= 1,494,837.2406 \pm 1050.0911 \text{ ng gRNA}. \end{aligned}$$

Here, the fittings of the mathematical model to the experimental data for the mean values of both optimised parameters and best vectors of parameters listed above are shown in red in Fig. S10(b) for the experimental replicates (upper panel) and their mean values (lower panel).

Figure S11 displays the evolution of the mean distances, here labeled as $D_m^-$ following the same procedure used in Fig. S7. Specifically, panel (a) shows how these distances decrease generation after generation of the MA in the fitting of the mixed infections at 20 ºC. The inset displays the evolution of the best distances at each improvement event. Panel (b) displays the same results for the competition experiments performed at 30 ºC.

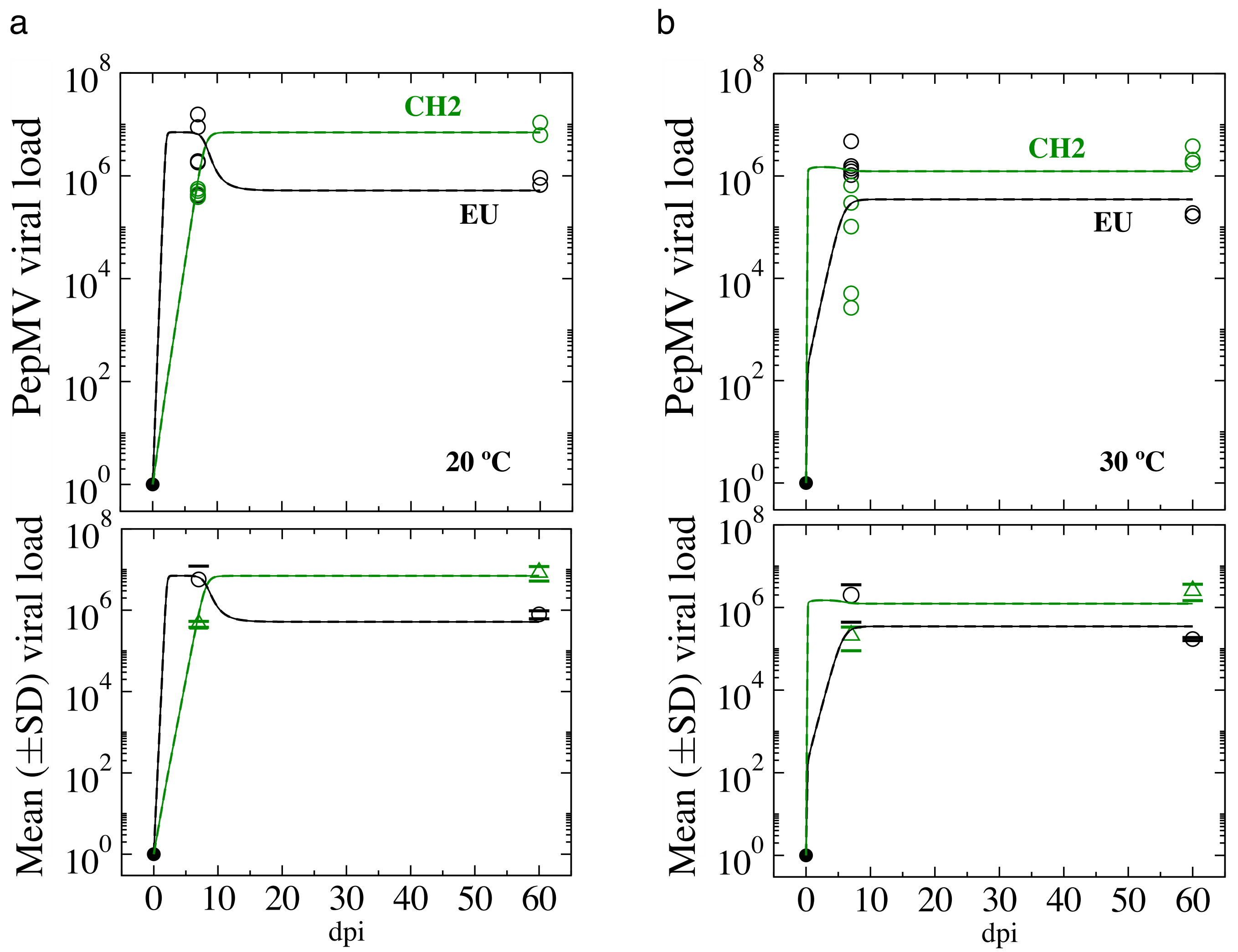

Figure S10: Fitting of the experimental data (open circles) for mixed infections (EU strain (black), CH2 strain (green)) using the mathematical model given by Eqs. (S.5)-(S.6). Panels (a) and (b) display the dynamics of competition at $T = 20^oC$ and $T = 30^oC$, respectively. Upper and lower plots display the viral load and the mean (± SD) viral loads, respectively. Here we show the fittings using the mean values of the optimised parameters (solid lines) and the mean values of the best parameter vectors (dashed lines). Note that since the values are extremely similar the time series appear overlapped.

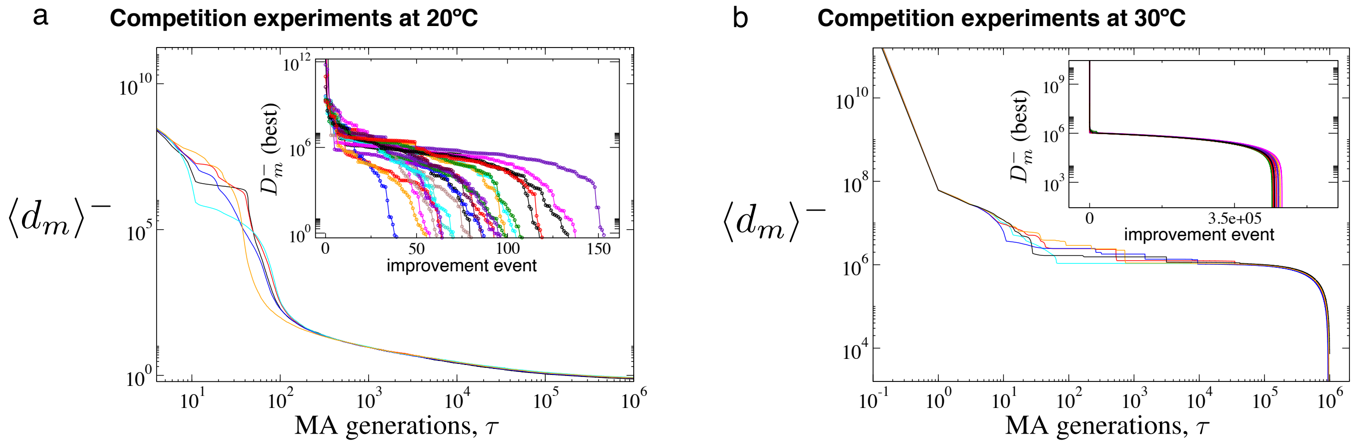

Figure S11: Same as in Fig. S7 for the fittings to the experimental data for mixed infections at 20ºC (a) and 30 ºC (b). The inset in (b) displays 10 replicates to ease visualisation since the curves appear overlaped.
